# An association between the gut microbiota and immune cell dynamics in humans

**DOI:** 10.1101/618256

**Authors:** Jonas Schluter, Jonathan U. Peled, Bradford P. Taylor, Melody Smith, Kate A. Markey, Ying Taur, Rene Niehus, Anna Staffas, Anqi Dai, Emily Fontana, Luigi A. Amoretti, Roberta J. Wright, Sejal Morjaria, Maly Fenelus, Melissa S. Pessin, Nelson J. Chao, Meagan Lew, Lauren Bohannon, Amy Bush, Anthony D. Sung, Tobias M. Hohl, Miguel-Angel Perales, Marcel R.M. van den Brink, Joao B. Xavier

## Abstract

The gut microbiota influences development and homeostasis of the mammalian immune system^1–3^, can alter immune cell compositions in mice^4–7^, and is associated with responses to immunotherapy that rely on the activity of peripheral immune cells^8–12^. Still, our understanding of how the microbiota modulates immune cells dynamics remains limited, particularly in humans where a lack of deliberate manipulations makes inference challenging. Here we study hundreds of hospitalized—and closely monitored—patients receiving hematopoietic cell transplantation as they recover from chemotherapy and stem cell engraftment. This aggressive treatment causes large shifts in both circulatory immune cell and microbiota populations, allowing the relationships between the two to be studied simultaneously. We analyzed daily changes in white blood cells from 2,235 patients, and 10,680 longitudinal microbiota samples to identify bacteria associated with those changes. Bayesian inference and validation across patient cohorts revealed consistent associations between gut bacteria and white blood cell dynamics in the context of immunomodulatory medications, clinical metadata and homeostatic feedbacks. We contrasted the potency of fermentatively active, obligate anaerobic bacteria with that of medications with known immunomodulatory mechanism to estimate the potential of the microbiota to influence peripheral immune cell dynamics. Our analysis establishes and quantifies the link between the gut microbiota and the human immune system, with implications for microbiota-driven modulation of immunity.

## MAIN TEXT

Experiments in mice provide evidence that the mammalian intestinal microbiome influences the development^1–3^ and homeostasis of its host’s immune system^4–7,13–15^. In humans, inflammatory bowel diseases correlate with functional dysbiosis in the gut microbiota^16,17^. Children born preterm and at term have different gut microbiome compositions and differ in the development of immune cell populations in their blood^18^. The composition of the gut microbiota may also influence the success of immunotherapies^8–11^. Immune checkpoint inhibitor therapy relies on activation of circulating T-cells and its success has, independently, been associated with abundances of intestinal anaerobic genera such as *Akkermansia*^9^ and *Faecalibacterium*^10^. It is therefore an intriguing prospect to augment treatments such as cancer immunotherapy^19^, including the burgeoning field of chimeric antigen receptor (CAR) T-cell therapy^20^, by leveraging microbiome-driven immune system modulation.

Our understanding of how the microbiota influences the dynamics of immune cells in humans, and how this compares to deliberate immunomodulatory interventions nevertheless remains limited. Experiments with animals may not always be sufficient to study mechanisms of microbiome-immune interactions and translate them to human biology as the microbial ecology in the gut of an animal model may be different from humans receiving treatment^21^. On the other hand, studies directly in patients may be criticized when they have small subject numbers, are cross-sectional, lack statistical power, or disregard key confounders such as medications^21^.

To overcome these limitations, we conducted a large-scale longitudinal study of the gut microbiota and day-by-day changes in circulatory immune cell counts. We investigated immune reconstitution dynamics after allogeneic hematopoietic cell therapy (HCT) within all 2,926 patients who underwent HCT at Memorial Sloan Kettering for various hematological malignancies, including leukemia, between 2003 and 2019 (**Figure 1A, Table S1**). The conditioning regimen of radiation and chemotherapy administered to HCT patients is the most severe perturbation to the immune system deliberately performed in humans and thus offers a unique opportunity to investigate dynamic links between the gut microbiota and the immune system directly in humans. Conditioning depletes white blood cell counts (**Figure 1A**) and can lead to prolonged periods of neutropenia (<500 neutrophils per *μ*l blood). Immune reconstitution begins after transplanted stem cells have matured sufficiently to release granulocytes from the bone marrow (neutrophil engraftment is defined as 3 consecutive days with >500 neutrophils per *μ*l blood, **Figure 1A-C**). The blood of each patient is carefully monitored throughout this recovery, and medications are administered to modulate the immune cell dynamics, including granulocyte-colony stimulating factor (GCSF) to increase neutrophil counts, and immunosuppressants such as mycophenolate mofetil or tacrolimus to prevent complications such as graft-vs-host disease (**Figure 1 J,K**). To investigate if the composition of the gut microbiota is associated with the dynamics of circulating white blood cells, we analyzed detailed blood and clinical metadata of our patients between 3 days before HCT and until 100 days post neutrophil engraftment (excluding pediatric patients, and other exclusion criteria: N=2,235, supplementary methods, **Figure S1**). During this period patients are monitored carefully, and our analysis included over 140,000 host phenotype measurements in the form of complete blood counts which quantify the most abundant white blood cells—neutrophils, lymphocytes, monocytes, eosinophils—as well as platelet counts (**Figure 1, S1**). We started collecting patients’ fecal microbiota data in 2009^22^, and by now obtained 10,680 high frequency, longitudinal microbiota compositions.

**Figure 1:**
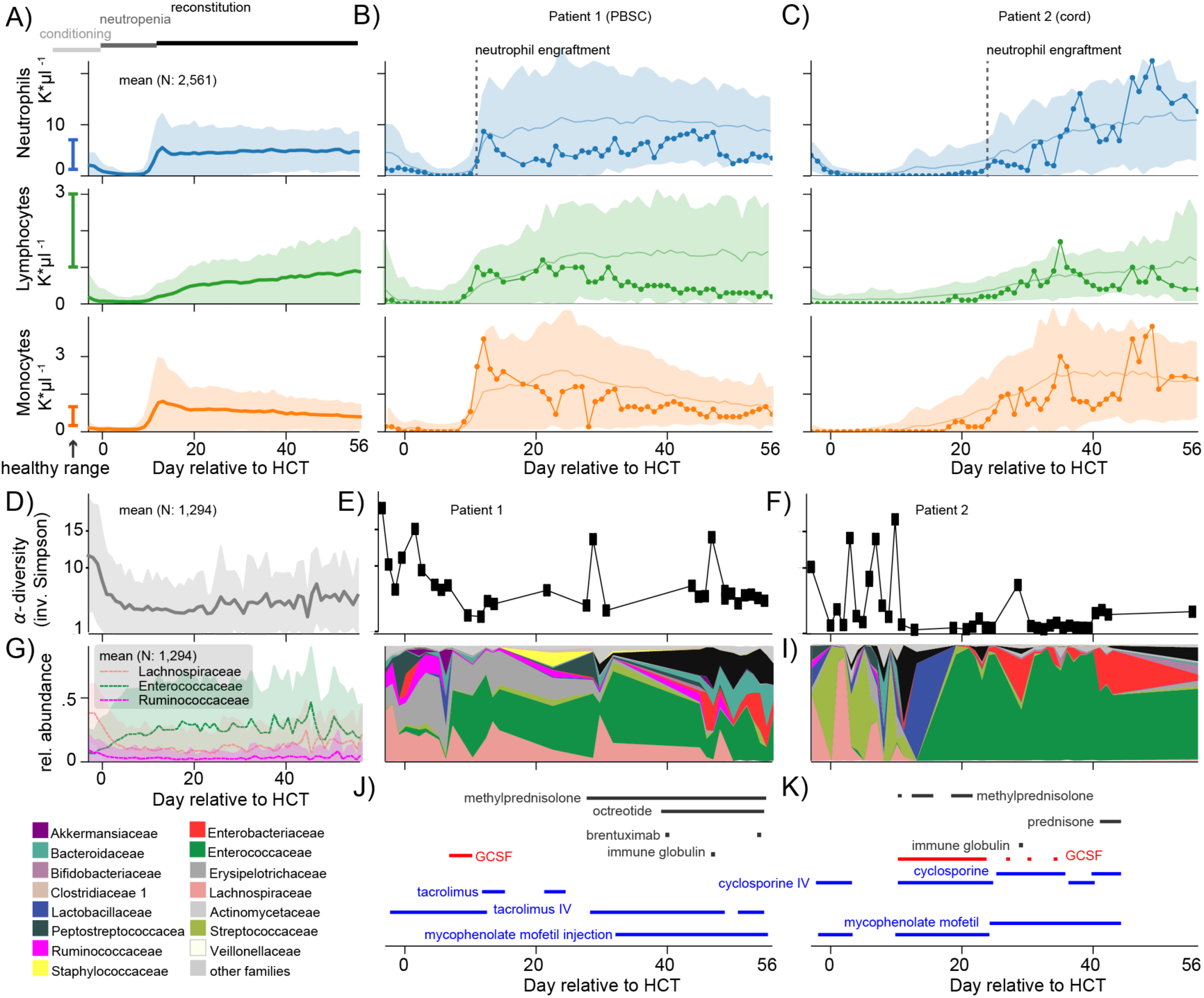
Immune reconstitution and microbiome dynamics after allogeneic hematopoietic cell transplantation (HCT). A) Three major periods of HCT—immunoablation during chemotherapeutic conditioning before HCT, defined as day 0, post-HCT neutropenia, and reconstitution following neutrophil engraftment—lead to recovery trajectories with large variability between patients. Shown are the mean counts (shaded: ±1 standard deviation, σ) of neutrophils, lymphocytes and monocytes per day relative to HCT from patients transplanted between 2003 and 2019 (A), contrasted with two individual patients (B,C) representative of the recovery trajectories for different stem cell graft source; patient 1 who received a peripheral blood stem cell graft, PBSC (line with circles: patient data, solid line and shaded region: mean±1 standard deviation of all PBSC patients), and patient 2 who received a graft of umbilical cord blood (line with circles: patient data, solid line and shaded region: mean±1 standard deviation of all cord patients). Fecal samples collected and analyzed by 16S rRNA gene sequencing reveal the loss of microbial diversity reported previously^23,24^ (D, line: mean per day, shaded: ±1 standard deviation); E,F: individual patient measurements) and commensal families (G, line: mean relative abundances of bacterial families, shaded: ±1 standard deviation); H,I: individual patient measurements), often replaced by Enterococcaceae domination. J,K). Administration of immunomodulatory medications for the two example patients.

HCT patients lose gut microbiota biodiversity and commensal microbial families during their treatment (**Figure 1D-I**); this figure generated from N=1,294 HCT patients clarifies the preliminary trends observed in previous studies from smaller datasets^23,24^. We have shown recently that mice with a depleted intestinal flora had worse recoveries of white blood cells after bone-marrow transplantation^25^. In our patients, microbial diversity usually recovers slowly during white blood cell reconstitution (**Figure 1D**); however, microbiota recovery as well as immune reconstitution can vary strongly between patients and treatment types (**Figure 1B,C,J,K, S1**). This variation is illustrated by the distinct trajectories of patient 1 who received a graft of peripheral blood stem cells (PBSC), retained high microbiota diversity and engrafted earlier (**Figure 1B,E,H**), and patient 2 who received a graft of umbilical cord blood (cord), lost microbiota diversity and engrafted later (**Figure 1C,F,I**). Low microbiota diversity at the time of neutrophil engraftment has been associated with 5-fold increased transplant-related mortality^26^, suggesting that that the joint recovery of the microbiota and white blood cells in circulation is critical for clinical outcomes.

To detect a directional and causal link between the microbiota and white blood cells, we first used data from a recent prospective randomized trial of autologous fecal microbiota transplantation (auto-FMT), which is a microbiota manipulation experiment done directly in our patients^23^ (supplementary methods). Twenty-four patients (**Figure 2A, Table S2)** underwent randomization, resulting in 10 untreated control and 14 treated patients, including patient 3 in **Figure S2**. To investigate if auto-FMT affected white blood cell reconstitution, we compared the 24 patients’ neutrophil, lymphocyte, monocyte (**Figure 2**) and total white blood cell counts (**Figure S3**) post-engraftment (i.e. when the transplanted hematopoietic cells begin producing new white blood cells, **Figure 2A**,**S3**). FMT procedures were conducted at variable time points relative to neutrophil engraftment, but overall, we observed higher counts of each white blood cell type in patients who received an auto-FMT during the first 100 days post neutrophil engraftment (p<0.001, **Figure 2B**,**S3**-**S6**).

**Figure 2:**
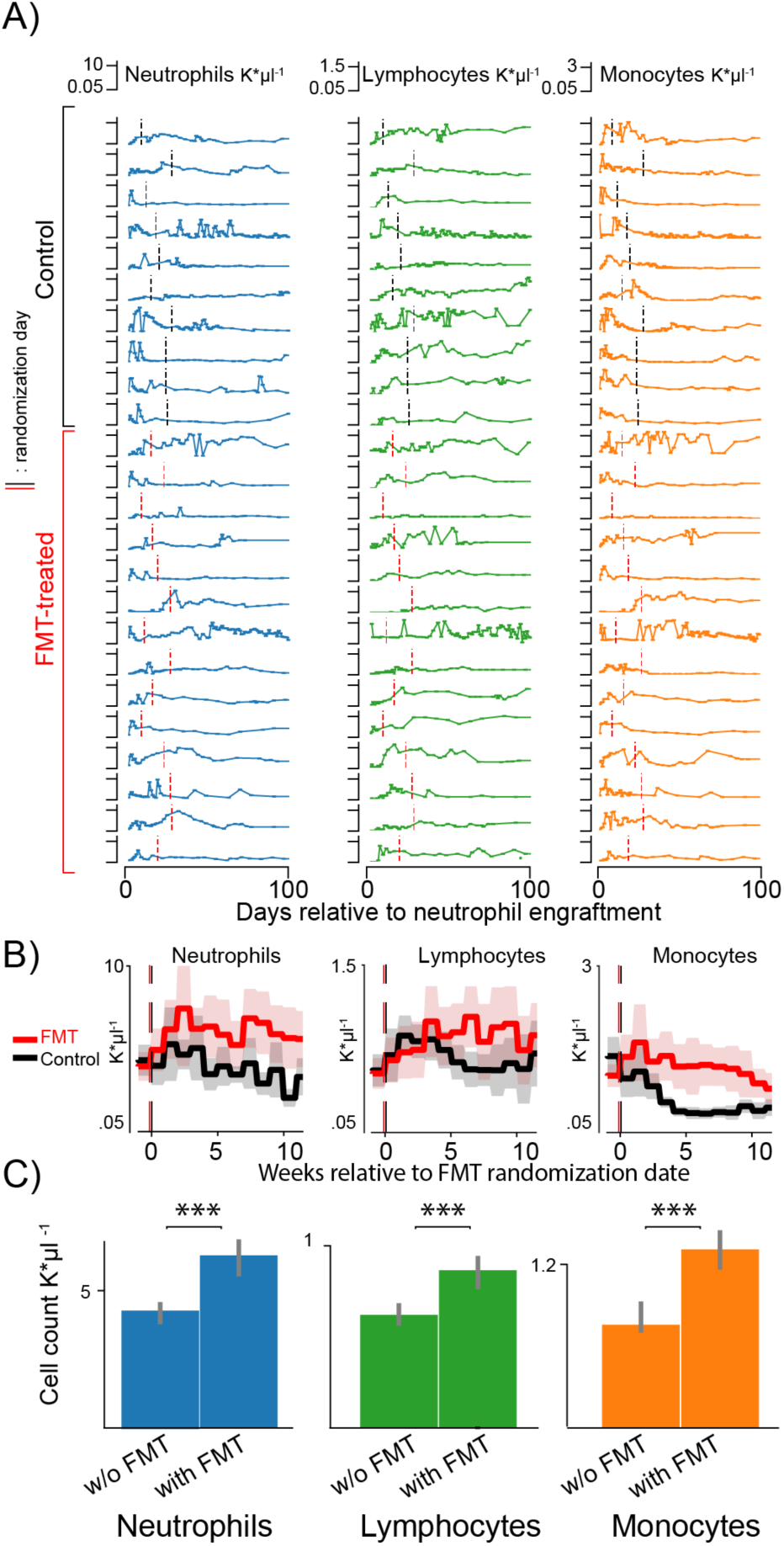
Neutrophil, lymphocyte and monocyte counts increased in FMT-treated patients during the weeks following the treatment compared to control patients. A) Absolute counts of neutrophils (blue), lymphocytes (green) and monocytes (orange) in 24 patients enrolled in a randomized controlled trial to receive an autologous fecal microbiota transplant post-neutrophil engraftment (10 control: black vertical line, 14 FMT treated: red vertical line,). B) Weekly mean cell counts aligned to the date of randomization into FMT treatment arm (red) or control (black). Line: mean per week, shaded region: 95%-CI. C) Results from a linear mixed effects model with random effects per patient and per day relative to neutrophil engraftment confirms that neutrophil, lymphocyte and monocyte counts are higher in patients receiving auto-FMT after their treatment as compared to control patients after the randomization date; bars and confidence intervals for the averages of observed white blood cell counts without FMT and post-FMT (***: p<0.001).

The increased white blood cells in patients receiving auto-FMT could be due to the reconstitution of a complex microbiota that we saw in these patients^23^ and the associated metabolic capabilities^6,7,25^, or it could be a systemic response to a severe therapy which introduced billions of intestinal organisms at once via an enema (no enema was administered to control patients^23^). Moreover, while the mixed-effects model accounted for patient-specific HCT treatments in this randomized patient cohort, chance differences in extrinsic factors such as different immunomodulatory drug exposures may have affected this result due to the small cohort size. Nonetheless, observing that auto-FMT recipients had increased white blood cell counts supports the notion that the microbiota can modulate the peripheral immune system. High counts of lymphocytes during immune reconstitution has been associated with improved clinical outcomes^27^. Additionally, in our HCT patients, a higher average level of white blood cells measured across a period of 100 days after neutrophil engraftment (supplementary methods) confirms a positive association with 3-year survival (hazard ratio: 0.91, p:0.04). Determining which taxa modulate immune dynamics could open new ways to improve robust immune reconstitution, which is critical for clinical outcomes^27–29^.

To address this question, we next investigated the link between the gut microbiota and the dynamics of white blood cell recovery in our large observational cohort of HCT patients. Homeostasis of circulatory white blood cell counts is a complex, dynamic process: neutrophils, lymphocytes and monocytes are formed and released into the blood *de novo* by differentiation of hematopoietic progenitor cells from the bone marrow, and they can be mobilized from thymus and lymph nodes (lymphocytes), spleen, liver and lungs (neutrophils); they can also migrate from the blood to other tissues when needed^30^. These processes are dynamic sources and sinks of circulatory white blood cells, and they can be modulated by drugs administered to patients receiving HCT. To identify factors associated with these dynamic source- and sink-processes—including the microbiota—we developed a two-stage approach analyzing the changes of white blood cell counts between two days (i.e. the rates of cell count increases and decreases). Stage 1 served as a feature selection stage where we used data of 1,096 patients (after filtering for qualifying samples and applying exclusion criteria, see supplementary methods) *without* available microbiome information to identify associations between clinical metadata, including immunomodulatory medications (methods), and changes of white blood cell counts from one day to the next (**Figure 3A**). Stage 2 was performed on data from an independent cohort of 841 different patients at MSK from whom concurrent microbiome samples were available. Stage 2, our main analysis, sought to reveal associations between the abundance of microbial taxa and the daily changes in blood cell counts in context of immunomodulatory medications, additional clinical metadata and the current state of the blood itself.

**Figure 3:**
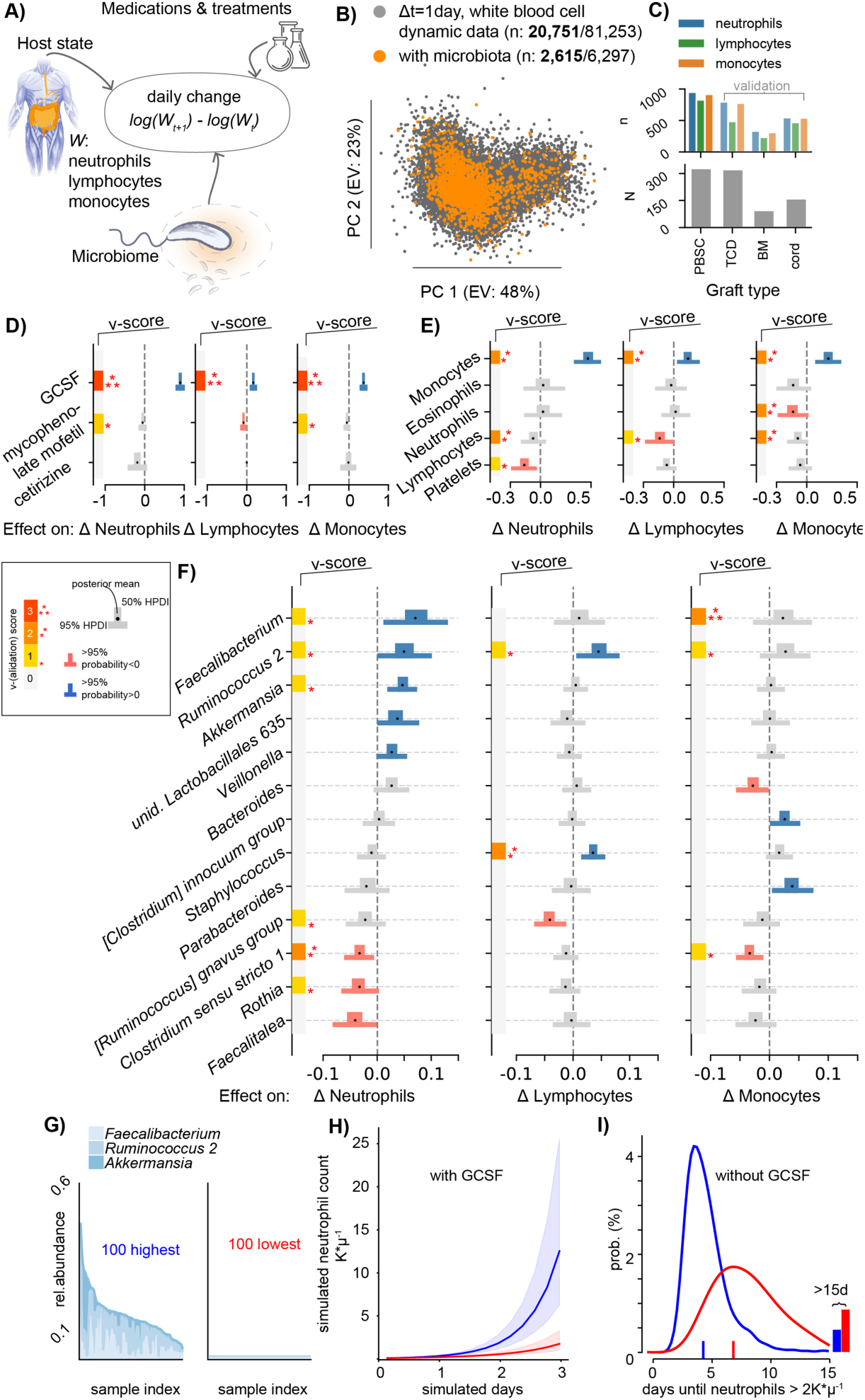
Bayesian inference conducted in context of the host state (white blood cell and platelet counts) and clinical variables including immunomodulatory medications reveals the microbiota potential to affect daily changes in circulatory white blood cell counts. A) Cartoon of the model: observed changes in white blood cell counts between two consecutive days are associated with the current state of the host in the form of blood cell counts in circulation, the administration of immunomodulatory medications, patient and clinical metadata, and the state of the microbiome. B) Visualization of the dynamic white blood cell data; scatter plot of the principal components (PC) of observed daily changes of neutrophils, lymphocytes and monocytes without (grey, see **Figures S7-9**) and with (orange) concurrent longitudinal microbiome data (bold: post engraftment sample counts). C) PBSC patients (N=312) provided the most blood samples with simultaneous microbiome data (n=995) relative to TCD, BM and cord patients, who were used as validation data sets. D-F) PBSC patient data inference results; bars show the posterior parameter estimate distributions (thin: 95% highest posterior density intervals, HPDI95, thick: HPDI50) for the jointly inferred associations between treatments (D), white blood cells counts (E), and fecal microbiota genus log-relative abundances (F) with the observed daily changes in neutrophils, lymphocytes and monocytes. v-score: number of validation cohort confirming associations, *always* set to zero if invalidated in any of the TCD, BM, or cord cohorts (additional coefficients in **Figure S10)**. G) 100 microbiota samples with highest (left) or lowest (right) relative abundances of *Faecalibacterium, Ruminococcus 2* and *Akkermansia*. H) Simulation of the neutrophil population over time in presence of GCSF with microbiota compositions sampled either from those high (blue) or low (red) in *Faecalibacterium, Ruminococcus 2* and *Akkermansia* relative abundance as shown in G); line: median of 1,000 simulations, shaded regions: quartile range of simulated neutrophil trajectories. I) In absence of GCSF, equivalent simulations to H) predict that the time to reach neutrophil counts >2K*μl^-1^ for the first time after HCT when the microbiota is high (red) in *Faecalibacterium, Ruminococcus 2* and *Akkermansia* compared with when these genera are low (blue) will decrease from 6.8 (95%-confidence interval, CI: [6.5, 7]) days to 4.4, (CI: [4.3, 4.5]).

In stage 1 we calculated the changes in neutrophil, lymphocyte and monocyte counts during patients’ recovery from >20,000 pairs of post-engraftment blood samples separated by a single day (**Figure 3B, S7-9**, supplementary methods). Using a cross-validated feature selection approach, we detected medications and HCT treatment parameters that were associated with different rates of change in neutrophil, lymphocyte and monocyte counts, including, as expected, GCSF and the graft stem cell sources (**Figure S7-S9, Table S1**).

During stage 2 we sought to identify associations between bacterial taxa of the gut microbiota and the dynamics of immune cells in circulation. For this, we performed Bayesian inferences using data from different sets of patients *with* available microbiome samples. Stage 1 had identified—as expected—that stem cell graft sources are associated with immune reconstitution kinetics (e.g. cord on average slower compared with PBSC^31^), and we therefore stratified our patients by graft source in stage 2. The model of stage 2 now included microbial genera as predictors of observed changes in white blood cell counts, in addition to the medications selected in stage 1, clinical features (conditioning intensity, age, sex), and the current state of the blood in the form of counts of neutrophils, lymphocytes, monocytes, eosinophils, and platelets. The data comprised 841 patients, but approximately 60% of the stool samples paired with a daily change in white blood cell counts were taken before neutrophil engraftment (**Figure 3B, Table S1**, supplementary methods), i.e. when blood cells counts were zero. In total, we analyzed 2,615 post-engraftment observations of changes in neutrophil counts during immune reconstitution (lymphocytes: 2,006, monocytes: 2,534) with paired stool samples which provided a large sample of observed white blood cell dynamics (**Figure 3B, Table S3**,**S4**). We first focused on the data from the largest (**Figure 3C**) cohort—patients who received a PBSC graft—and withheld the other cohorts (bone marrow, BM; T-cell depleted graft (*ex-vivo*) by CD34+selection, TCD; and cord) to use as independent validation cohorts. For this validation, we analyzed TCD, BM, and cord patients’ data in the same way as PBSC patients’ data and compared the resulting posterior coefficient distributions (methods). We assigned coefficients obtained from the PBSC cohort a validation score (v-score) between 0 and 3, representing the number of times that the focal coefficient was validated in the other cohorts; but, conservatively, the score was always set to zero if we observed counter-evidence among any of the other data sets, i.e. evidence that coefficients had the opposite sign, ensuring only the most consistent associations were considered as validated. Finally, we analyzed data from another patient cohort consisting of 493 bone marrow transplantation recipients treated at Duke University including 9,603 blood samples and a total of 629 microbiota samples from 218 patients, albeit with lower sampling density, and we used the results from this analysis for further validation.

Notably, as a verification of our approach, we detected associations between the administration of immunomodulators and increased or decreased rates of immune cell count changes consistent with the known biological mechanism of these medications (**Figure 3C**,**S10-S13**). The strongest across all predictors is the well-known neutrophil-increasing effect of GCSF^32^; GCSF administration—used to accelerate recovery from chemotherapy-induced neutropenia^32^—was associated with a +140% increase in the rate of neutrophil changes from one day to the next ([+114%, +170%], 95 percent probability density interval [HPDI95]). This finding was observed in all MSK validation data sets (v-score=3, **Figure 3D**), as well as among Duke University patients (**Figure S14**,**S15**). We furthermore found a GCSF-associated increase of +43% ([+30%, +58%]HPDI95, v-score=3) in monocyte rates, and, although smaller, in lymphocyte rates (+16%, [+5%, +27%]HPDI95, v-score=3). Both neutrophil and lymphocyte rates decreased following the exposure to antihistamine or immunosuppressive medications (cetirizine -18%, [-35%, +5%]HPDI95, mycophenolate mofetil -8% [-15%,+1%]HPDI95, respectively). Finally, less intensive chemotherapeutic conditioning regimens (non-ablative conditioning and reduced intensity) were associated with larger lymphocyte and monocyte count growth rates during immune reconstitution (**Figure S10C**)

Beyond associating medications in agreement with their known biological mechanism, our analysis detected associations between the current count of white blood cells and their rate of change: a negative association among lymphocytes, negative associations between counts of neutrophils and lymphocytes with the rates of monocytes, and a negative association between the counts of platelets and lymphocytes and the rates of neutrophils (**Figure 3E**). Conversely, we found positive associations between monocytes and the rates of each of the investigated white blood cell subsets. These associations, derived from daily counts of white blood cells, could reflect a complex network underlying the regulation of blood immune cell composition^30^. More importantly, the associations quantified for medications and potential homeostatic feedbacks provided a benchmark against which we could compare associations from gut microbial taxa.

We identified microbial genera that consistently associated with increases or decreases in white blood cell counts by first using data from the PBSC patients and then validating the associations in the other cohorts (**Figure 3F**). Higher abundances of *Faecalibacterium* (+8%, [+1%, +14%]HPDI95 per log_10_), *Ruminococcus 2* (+5%, [0%, +10%]HPDI95) and *Akkermansia* (+4%, [+1%, +7%]HPDI95) were associated with greater neutrophil increases, whereas increased *Rothia* (- 3%, [-7%, 0%]HPDI95), and *Clostridium sensu stricto 1* (−3%, [-6%, 0%]HPDI95) relative abundances associated with reduced neutrophil rates. These results were validated in univariate analyses conducted in the Duke University cohort (**Figure S14, S15**). We also conducted the inference using total genus abundances as predictors instead of relative abundances; this analysis confirmed *Faecalibacterium* as most strongly associated with neutrophil dynamics (**Figure S16**, supplementary methods). *Staphylococcus* was positively associated with lymphocyte rates (+4%, [+1%, +6%]HPDI95) and, again, *Ruminococcus 2* was also associated with faster lymphocyte increases (+5%, [+1%, +9%]HPDI95). Both *Faecalibacterium* as well as *Ruminococcus 2* also associated with increases in monocytes, and while this association was validated in other cohorts (v-score 3 and 1, respectively), there was higher uncertainty of the association estimate (HPDI50>0). Again, *Clostridium sensu strictu 1* (−3% [-5%, -1%]HPDI95) associated consistently with decreased rates of monocytes. The associations we identified—and validated in other cohorts—between microbial taxa in the gut and daily changes in white blood cell counts support the idea that hematopoiesis and mobilization respond to the composition of the gut microbiome, influencing systemic immunity^33^.

Most of the taxa that strongly associated with white blood cell dynamics were obligate anaerobes. *Rothia*, was a notable exception: this aerobic genus is typically found in the oral cavity^34^ but can become an opportunistic pathogen in immunosuppressed patients and is not known to provide metabolic functions to the host^35^. Some obligate anaerobes, on the other hand, produce short-chain fatty acids^36,37^ and bacterial cell-wall molecules^1,38,39^ that modulate immune responses and granulopoiesis^6^. Nutritional support from the intestinal microbiota improved hematopoietic reconstitution in a mouse model^25^. To identify a similar association in our patients, we estimated a microbiota potency by multiplying the log_10_-relative abundances of microbial genera in a sample with their corresponding posterior coefficients. We analyzed shotgun metagenomics sequences from 124 of the samples and observed that samples with positive microbiota potency were associated with enrichment in cholate degradation and vitamin-B1 synthesis related pathways, as well as butanoate formation (**Figure S17**). Our findings are in line with evolutionary theory^40^ that essential but broadly available microbial traits such as the production of B vitamins, secondary bile acid metabolism, and fermentation to short-chain fatty acids^41^ could be co-opted by the host’s immune system as part of the homeostatic interplay between immune system and a complex microbiota^42,43^. For example, *Ruminococcus 2* is a genus that contains *R. bromii*, a keystone species necessary for the efficient release of energy from complex starch in the normal diet^44^. Reassuringly, the genera *Faecalibacterium, Ruminococcus 2* and *Akkermansia* that we associated with faster rates of white blood cells (**Figure 3F**) were among those best reconstituted by auto-FMT^23^, potentially explaining why we found higher counts of neutrophils, monocytes and lymphocytes in patients who received the auto-FMT (**Figure 2B,C**).

The associations we reveal are interpretable as potential effectors on sources and sinks of white blood cell counts in circulation. Intestinal bacteria may affect white blood cell counts in circulation by influencing either their sources in the bone marrow or their cytokine profiles^45^ and proliferation rates in the blood, their sinks in different organs, or both. The human immune system in turn can interact with the microbiota and modulate its composition, for example via immunoglobulin A responses targeting specific bacteria as studied in mice^43,46,47^. To investigate a reverse effect of the peripheral immune cell system onto bacterial populations, we employed an analogous approach to the stage 1 analysis of white blood cell dynamics. Dynamics of white blood cells can be estimated from changes in absolute cell counts, and to obtain the necessary measurements in absolute bacterial abundances, we measured total bacterial 16S rRNA gene copies per gram of stool for a subset of our samples (3,995 samples from 481 patients). Using absolute abundances of bacteria as predictors in addition to medications, we jointly inferred the association network of dynamics between the gut bacterial ecosystem and the peripheral immune system. All of our patients receive antibiotics on some days during their treatment^24^ and their strong effects on microbiota dynamics were the dominant effects that survived cross-validated regularized elastic net regression (**Figure S18**). Relaxing the strength of the regularization (methods), however, revealed several bi-directional relationships between immune cells in circulation and bacterial dynamics in the gut (**Figure S19**). Of note, we detected a negative association of absolute *[Ruminococcus] gnavus* group abundance with lymphocytes dynamics, confirming our main result based on relative bacterial abundances (**Figure 3**). In the reverse direction, we saw a positive association of *[Ruminococcus] gnavus* group dynamics with lymphocyte counts. This result agrees with findings that *Ruminococcus gnavus* thrives in and promotes inflammatory conditions such as Crohn’s disease and other inflammatory bowel diseases (IBD)^48^; our analysis suggests it may drive high neutrophil to lymphocyte ratios that are broadly characteristic for poor disease outcomes in IBD^49,50^ and beyond^51,52^.

Overall, our analysis identified that the microbiome is associated with immune cell dynamics in addition to medications. The effects should be interpreted as net effects since they do not distinguish, for example, how the microbiota impacts *de novo* hematopoiesis in isolation from its impact on other sources and sinks. Unlike the plausible role of obligate anaerobe fermenters in augmenting hematopoiesis via nutritional support^25^, the positive association detected between *Staphylococcus* and lymphocyte dynamics could instead result from reduced extravasation of T cells from circulation into the gut epithelium^53^, especially since high abundances of *Staphylococcus* are associated with low gut microbiota diversity (p<0.001, **Figure S20**), which indicates a depleted microbiota.

Nevertheless, our approach allows us to leverage the chronology of events and assess “mathematical causality”^54^. Of course, due to the observational nature of these data there are risks of confounding that could explain some of the associations found, but the close temporal correspondence^54^ between microbiota and blood cell dynamics, and the validation across cohorts reduces the number of plausible confounders. Our results, therefore, quite naturally suggest candidate microbial taxa to manipulate if we seek to steer complex hematopoietic dynamics and utilize the microbiota as an immunomodulatory component of the human body. Intriguingly, members of *Faecalibacterium* and *Ruminococcus* in one study^10^, and *Akkermansia* in another^9^, were identified as enriched in patients with better responses to anti–PD-1 immunotherapy, which suggested a disagreement between the two studies^55^. Our results, however, revealed *Faecalibacterium, Ruminococcus 2*, and *Akkermansia* as the most strongly associated taxa with increases in white blood cell counts from one day to the next. Therefore, our results agree with the findings of both anti PD-1 therapy studies that these taxa are associated with immune modulation in humans. Our results also allow us to compare the potency of manipulating these intestinal commensals to that of immunomodulatory drugs. While these genera are common in the gut microbiota of healthy people^17^, the relative abundance of each genus can drop below detection in our patients during the intestinal damage related to HCT^24^. Therefore, realistic ranges of 3-5 orders of magnitude in bacterial log-relative abundances (**Figure 3G, S21**) can yield effect sizes similar to that of homeostatic feedbacks between white blood cells and several immunomodulatory medications (e.g. a change in *Ruminococcus 2* from below detection to 1% relative abundance associated with a +67% and +63% increases in neutrophil and lymphocyte rates, respectively). Therefore, while the effect sizes of intestinal bacteria at first may appear smaller than those of immunomodulatory drugs, the herein estimated homeostatic effects of gut bacteria may not be that small since their coefficients refer to changes in exponential rates of white blood cells and accumulate each day. To better demonstrate how this accumulation of effects would work, we conducted simulations of the inferred dynamic system of white blood cells using our posterior coefficient distributions (methods). We simulated 1,000 time series for microbiota compositions either chosen from the 100 samples highest or lowest in *Faecalibacterium, Ruminococcus 2* and *Akkermansia* (**Figure 3G**), in presence (**Figure 3H**) or absence (**Figure 3I**) of GCSF administration. Simulations predict that a microbiota enriched in these genera could accelerate immune reconstitution, and reduce the time until neutrophils reach >2K\**μ*l^-1^ in absence of GCSF by 2.4 days, from predicted 6.8 (CI: [6.5, 7]) to 4.4 days (CI: [4.3,4.5]) days. Gut bacteria, in concert and over time, could therefore have significant impact on systemic immunity even in individuals with less severely injured microbiomes.

In sum, our work links the gut microbiota to the dynamics of the human immune system via peripheral white blood cell populations. Our analysis uses white blood cells counted directly from patients, which are coarse-grained clinical analyses conducted at large scale but lack details such as lymphocyte and other immune cell subsets. Nonetheless, because it is in humans, this study fills an important gap at a critical time for microbiome research when the clinical relevance of animal models of microbiome-immune interaction has been questioned^21^. By studying a large number of patients over time, we were able to infer and quantify for the first time the association of microbiota components on systemic immune cell dynamics, and our results help to consolidate previous findings^10,9^ that seemed in conflict with each other^55^. Our study demonstrates that the composition of the microbiota does indeed modulate systemic immune cell dynamics, a link that could be used in the future to improve immunotherapy and help identify microbiota treatments for inflammatory diseases^9,10,56–60^.

## Supplementary Information

Appended and available online.

## Acknowledgments

We thank Marc Lipsitch, Sandra B. Andersen, Kevin R. Foster, Jonathan Kevin Sia, Eric G. Pamer, Kat Coyte, Sibylle Mitschka and the members of the Xavier lab for helpful discussion and comments on the manuscript. This work was supported by the National Institutes of Health (NIH) grant U01 AI124275 to JBX and grant R01 AI137269 to JBX, by the MSKCC Cancer Center Core Grant P30 CA008748, the Parker Institute for Cancer Immunotherapy at Memorial Sloan Kettering Cancer Center, the Sawiris Foundation, the Society of Memorial Sloan Kettering Cancer Center, MSKCC Cancer Systems Immunology Pilot Grant and Empire Clinical Research Investigator Program. MS received funding from the Burroughs Wellcome Fund Postdoctoral Enrichment Program, the Damon Runyon Physician-Scientist Award, and the Robert Wood Johnson Foundation. MRMvdB and JUP received financial support from Seres Therapeutics. TMH is investigator in the Pathogenesis of Infectious Diseases from the Burroughs Wellcome Fund, and funded via an award from Geoffrey Beene Foundation, and NIH RO1 AI093808. M-AP has received honoraria from AbbVie, Bellicum, Bristol-Myers Squibb, Incyte, Merck, Novartis, Nektar Therapeutics, and Takeda; has received research support for clinical trials from Incyte, Kite (Gilead) and Miltenyi Biotec; and serves on data and safety monitoring boards for Servier and Medigene and scientific advisory boards for MolMed and NexImmune. The funders had no role in study design, data collection and analysis, decision to publish, or preparation of the manuscript.

## Author Contributions

J.S. and J.B.X. wrote the manuscript. J.S. and J.B.X. designed the analyses with expert help from R.N.. J.U.P. and Y.T. contributed to the clinical data preparation, B.P.T. provided the 16S data processing pipelines, A.D. provided the shotgun processing pipelines. All authors contributed to the writing and interpretation of the results.

## SUPPLEMENTARY INFORMATION

### Methods

#### Complete blood count collection and characterization

Absolute white blood cells count data were obtained from routine complete blood counts ordered by clinicians during normal clinical practice, used to obtain informative diagnostic and monitoring information. Blood samples received in the clinical hematology laboratory were analyzed using Sysmex XN automated hematology analyzers (Sysmex, Lincolnshire, IL) and, when needed based on specific flags and parameters as per MSKCC standard operating procedures, were validated manually using the Sysmex DI-60 Slide Processing System or CellaVision DM9600 Automated Digital Morphology System (Sysmex, Lincolnshire, IL).

#### 16S rRNA gene amplification and multiparallel sequencing

For each sample, duplicate 50-μl PCRs were performed, each containing 50 ng of purified DNA, 0.2 mM deoxynucleotide triphosphates, 1.5 mM MgCl2, 2.5 U Platinum Taq DNA polymerase, 2.5 μl of 10× PCR buffer, and 0.5 μM of each primer designed to amplify the V4-V5: 563F (5′-nnnnnnnn- NNNNNNNNNNNN-AYTGGGYDTAAAGNG-3′) and 926R (5′-nnnnnnnn-NNNNNNNNNNNN-CCGTCAATTYHTTTRAGT-3′). A unique 12-base Golay barcode (Ns) precedes the primers for sample identification^61^, and one to eight additional nucleotides were placed in front of the barcode to offset the sequencing of the primers. Cycling conditions were 94°C for 3 min, followed by 27 cycles of 94°C for 50 s, 51°C for 30 s, and 72°C for 1 min. For the final elongation step, 72°C for 5 min was used. Replicate PCRs were pooled, and amplicons were purified using the QIAquick PCR Purification Kit (Qiagen). PCR products were quantified and pooled at equimolar amounts before Illumina barcodes and adaptors were ligated, using the Illumina TruSeq Sample Preparation protocol. The completed library was sequenced on an Illumina MiSeq platform following the Illumina recommended procedures with a paired-end 250 × 250 bp kit

#### Sequence analysis

The 16S (V4-V5) paired-end reads were merged and demultiplexed. Amplicon sequence variants (ASVs) were identified using the Divisive Amplicon Denoising Algorithm (DADA2) pipeline including filtering and trimming of the reads^62^. Reads were trimmed to the first 180 bp or the first point with a quality score Q<2. Reads were removed if they contained ambiguous nucleotides (N) or if two or more errors were expected based on the quality of the trimmed read. We assigned taxonomy to ASVs using a 8-mer based classifier trained by IDTaxa^63^ using the SILVA database^64^. We determined the copy number of 16S rRNA genes per gram of stool for 4,158 of our samples as reported previously^24^, by quantitative PCR on total DNA extracted from fecal samples.

#### Quantification of total microbiota density per gram of stool and estimation of total genus abundances

Quantitative PCR (qPCR) was performed on DNA extracted from the 1g wet weight of a stool sample using DyNAmo SYBR Green qPCR kit (Finnzymes) and 0.2 μM of the universal bacterial primer 8F (5′-AGAGTTTGATCCTGGCTCAG) and the broad-range bacterial primer 338R (5′-TGCTGCCTCCCGTAGGAGT-3′). Standard curves were prepared by serial dilution of the PCR blunt vector (Invitrogen) containing 1 copy of the 16s rRNA gene. Cycling conditions were 95°C for 10 minutes followed by 40 cycles of 95°C for 30 seconds, 52°C for 30 seconds, and 72°C for 1 minute. We used the measurements of total 16S rRNA gene counts per gram of stool to multiply the relative abundances of taxa obtained from 16S amplicon sequencing to obtain the estimate of their total abundance per gram of stool (supplementary methods). Importantly, this does not account for 16S copy number variation between taxa, but the observed dynamic ranges in total abundances of taxa in our data set span up to 9 orders of magnitude, exceeding the potential inaccuracies due to copy number variation.

#### Diversity calculations

Microbiome alpha-diversity was measured by the inverse Simpson (IS) index of a sample. It was calculated by 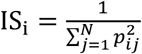, where *p* is the relative abundance of the *j*th ASV out of *N* total ASVs in sample *i*.

#### Linear mixed-effects model of white blood cell counts

To study the effect of auto-FMT on white blood cells, we investigated the white blood cell counts of 24 enrolled patients of this trial from the day of neutrophil engraftment until 100 days after. FMT occurred on different days relative to neutrophil engraftment. Thus, we performed an analogous analysis to that conducted in the original publication that demonstrated how FMT re-established a diverse microbiome in the post-FMT period^23^. To answer if white blood cell counts differed post-FMT, we used a linear mixed effects model of white blood cell counts, *y*, modeled as a function of the FMT treatment as well as patient and timepoint specific random effects. We included random intercept terms for each day *i* and each patient *j*, and a fixed effects term for the post-FMT period with associated coefficient “armpost”, using the indicator variable “FMT”, that is 1 when a patient was from the FMT treated arm of the trial and *day* was greater than or equal to the day of the FMT procedure. We conducted independent analyses for neutrophil, lymphocyte and monocyte counts. This resulted in the following model of a cell count, *y*, for patient *j* on day *i*:

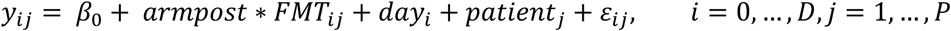

with prior distributions 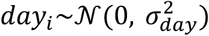, and 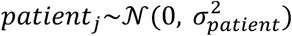, independent error *ε*_*ij*_∼*𝒩*(0, *σ*^2^) and fixed intercept β_0_, for the D days post neutrophils engraftment and P patients, (D=100, P=24). For convenience of those interested in reanalyzing our data, the part of our data concerning the auto-FMT analysis is available in tidy format (supplementary data 11), and the analysis code conducted in the R programming language is available as an exported notebook (fmt_effect_on_wbc.pdf) on github: https://github.com/jsevo/wbcdynamics_microbiome/.^65^ We conducted an additional analysis with “day” as a continuous predictor which did not change our conclusions (supplementary methods).

#### Dynamic systems analyses

We analyzed factors associated with the observed changes of absolute counts of neutrophils, lymphocytes and monocytes between two days. In the following we describe how chronology of events and biological samples were encoded, and the models used to infer a role of medications, clinical parameters and the microbiome on dynamics of white blood cells.

To reveal factors that associate with day-to-day changes in white blood cell counts, we started from a first-order differential equation of white blood cell (W) dynamics:

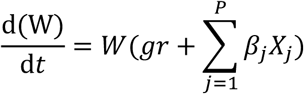

Where g*r* represents the intercept, i.e. the base line rate of change during immune reconstitution, and *β*_*j*_ are the to-be-estimated coefficients of the *P* predictors *X*_*j*_, *j* ϵ *P*, of the white blood cell dynamics.

This equation was then linearized to

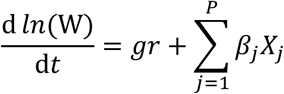

And we parameterized the corresponding discrete difference equation:

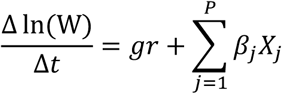

where Δln(W) is the log-difference between single days of neutrophils, lymphocytes or monocytes counts, and Δt=1 for all intervals. Predictors include the counts of neutrophils, lymphocytes, monocytes, eosinophils and platelets during an interval (homeostatic feedbacks), immunomodulatory medication and clinical observations such as a blood stream infection and the onset of graft versus host disease, HCT parameters such as graft types and conditioning regimens, and, additionally, the microbiota composition in “stage 2” of our analysis (supplementary methods for data exclusion and additional details on interval definitions). Importantly, by parameterizing a dynamic equation and analyzing *rates of change*, our coefficient estimates have an immediate causal interpretation within our modeling framework (i.e. a *β*_*j*_>0 implies that higher levels of the corresponding *X*_*j*_ *increases* the respective white blood cell type, *W*). To differentiate such results from other associations, analyses of this type have been termed “mathematical causality”^54^.

##### Stage 1 analysis: Feature selection. Identifying medications and clinical observations associated with white blood cell dynamics from patients without microbiome data

Stage 1 uses data of patients without any available microbiome samples and the following model of white blood cell changes, y:

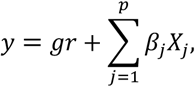

with intercept, *gr*.The predictors, *X*, include dummy variables for the HCT graft type, patients’ age on the date of HCT, sex, 13 most frequently observed positive blood cultures with remaining other blood stream infections grouped into a separate category “other infections”, an indicator for the onset of graft versus host disease, administrations of 55 different, most common immunomodulatory medications and platelet transfusion events, and HCT conditioning intensity regimens as well as the log-transformed geometric mean counts of neutrophils, lymphocytes, monocytes, eosinophils and platelets during the respective interval. We used elastic net regression^66^ for feature selection using the sklearn package for the Python programming language^67^. For elastic net regression with 50% L1- penalty, predictors were scaled between zero and 1, and we used 10-fold cross validation (i.e. leaving out 10% of patients at each cross-validation step) to choose the regularization strength, *λ*, solving for

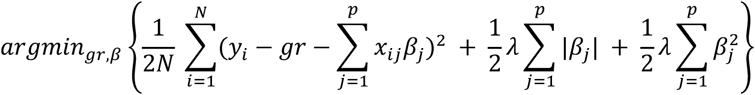

Stage 1 yielded a sparse coefficient matrix of predictors used to design the model in stage 2.

##### Expanded analysis on patients with microbiome data – stage 2

To identify associations between microbiota and white blood cell dynamics, we conducted an analogous, Bayesian regression using the package PyMC3 for the Python programming language^68^. Stage 1 identified important difference between transplant types, and we therefore stratified our data into 4 cohorts according to their stem cell graft source. Using data independently from each cohort, swe applied “no U-turn” sampling^69^ to produce 10,000 posterior samples from 5 independent MCMC chains that parameterized the model:

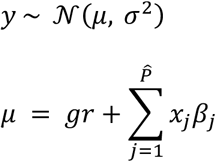

with uninformative prior distributions

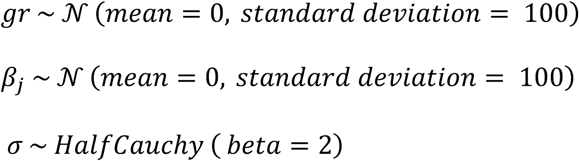

where *y* is the observed daily change of a focal white blood cell type as in stage 1 with normal distributed mean, *μ*, and *σ*, the model uncertainty with a thick-tailed half Cauchy prior (importantly, our posterior estimates do not depend on this choice as we obtain the same results with an inverse Gamma prior, figure S19). *μ* was a function of the baseline growth rate, *gr*, and predictors, 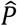: medications with non-zero coefficients in stage 1, the white blood cell counts, patient age and sex, and HCT conditioning intensities; additionally, 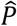 now included the log-abundances of microbial genera as measured by 16S sequencing from DNA in the stool collected on the second day of a daily interval (see supplementary methods for details). We considered taxa that were among the 100 most abundant, or had reached maximum relative abundances of at least 10%, and selected those who were non-zero in more than 75% of our samples. White blood cell counts and microbiota data present during a daily interval were log-transformed, and zeros were filled with half of the minimum observed non zero counts (i.e. 0.5e3 and 2e-6, respectively). We focused on the largest cohort (PBSC) and used the independent inference results from TCD, BM, and cord cohorts for validation.

##### Validation score

Coefficients learnt from the PBSC patient cohort were assigned a “validation score” based on the results obtained from the other three MSK patient cohorts. Our requirements for validation were conservative; we required evidence from our validation data sets as well as absence of counter evidence. For regression results from each of the validation graft type cohorts, i.e. TCD, BM, and cord, we checked if a coefficient had more than 75% probability (50%HPDI) to have the same sign as the mean of the PBSC coefficient posterior for a given predictor. If so, this was considered evidence of validation, and we summed the evidence over the three validation sets (i.e. maximum score of 3, 1 from each of TCD, BM, and cord cohorts). Conversely, if we found more than 75% probability among any of the validation data sets that a given predictor had the *opposite* sign as the posterior mean calculated from PBSC data, this was considered counter evidence and the validation score was always set to zero.

##### Analysis of white blood cell dynamics with absolute bacterial abundances as predictors instead of relative abundances

We conducted an ordinary least squares regression using the statsmodels package in the Python programming language of the same model as in the main Bayesian analysis using total bacterial abundances as predictors. This was only possible on a subset of 389 neutrophil, 331 lymphocyte and 376 monocyte rate observations from PBSC patients.

##### Forwards simulation of predicted immune system reconstitution kinetics

To assess the impact of the estimated microbiota coefficients on immune system dynamics, we conducted 1,000 simulations of the system of 3 differential equations describing the dynamics of neutrophils, lymphocytes and monocytes. We ran 1,000 simulations four times: in presence and absence of GCSF, each with microbiota compositions enriched or depleted in *Faecalibacterium, Ruminococcus 2* and *Akkermansia*. To identify these compositions, we ranked the observed microbiota compositions by these taxa, and chose randomly either from the top or bottom 100. The coefficients for white blood cell interactions, interactions with the microbiota and the effect of GCSF were sampled from our posterior coefficient distributions. Using these coefficients sampled at the start of the simulation, and using 50 cells*μl^-1^ of neutrophils, lymphocytes and monocytes as initial values, we simulated these differential equations forwards in time using the odeint function of the scipy package for the Python programming language.

##### Validation on data from Duke University

We analyzed 9,603 blood samples with 25,581 associated administrations of immunomodulatory medications, and 741 microbiota samples from Duke as an orthogonal data set to validate our findings. The temporal resolution of this data was much lower, and after filtering for samples from the relevant post neutrophil engraftment period, and by requiring daily intervals, 83 valid, complete data points were available. Using these data, we correlated daily blood cell changes individually in univariate, or jointly in a partial least squares regression, with those predictors that achieved more than 95% probability density in the positive or negative domain in the PBSC data regression. For each of these predictors, we present the sign of slopes and Bonferroni corrected *p*-values from individual linear regressions.

##### Joint analysis of the effect of antibiotics and white blood cell counts on the microbiota and the microbiota and immunomodulatory medications on white blood cell counts

Analogous to *stage 1*, we performed cross-validated, regularized linear regressions (ElasticNet) using the scikit-learn package for the Python programming language to jointly estimate the association network between microbiota and circulatory white blood cells. For this, we constructed a block matrix **X** of predictor matrices ***X***_***i***_ that include the absolute bacterial abundances, drug data (antibiotics for bacterial dynamics and immune modulators for white blood cell dynamics), as well as the counts of white blood cells and a separate intercept term per block. Each block 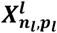, with *n*_*l*_ observations and *p*_*l*_ predictors (l=0…k), on the diagonal of **X** corresponds to the indices of the observed daily log-changes of one of the 41 bacterial genera considered in our main analysis or the log changes in neutrophil, lymphocyte and monocyte counts from PBSC patients contained in **Y** (in total we calculated 15,833 rates from 256 patients). Our regression problem can thus be written as:

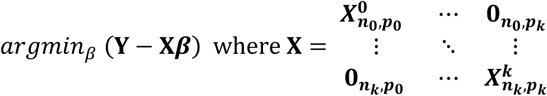

with k=44, i.e. 41 bacterial genera and 3 white blood cell types, the to-be estimated coefficient vector **β** and **0** the zero matrix. This system is underdetermined and we therefore chose the same approach as in stage 1, elastic net regression, for feature selection. Predictors were scaled between zero and 1, and we used 3-fold cross validation, leaving out 1/3^rd^ of the patients at each iteration to identify a global regularization strength, *λ*, solving for

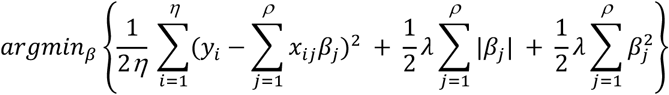

where *η* is the total number of observed daily log changes in genera and white blood cells, and *ρ* the total number of predictors. This yielded a strongly regularizing *λ*_*s*_, and thus few predictors. To characterize potential bidirectional relationships between white blood cell counts and the gut microbiota, we iteratively reduced the regularization strength until the strongest interaction between microbiota and white blood cell dynamics, i.e. *Faecalibacterium* with neutrophil dynamics, was detected. We than re-ran the regression with this reduced regularization strength, *λ*_*r*_.

##### Shotgun sequencing

Sequencing of 124 post-neutrophil engraftment was conducted on the Illumina HiSeq platform. For details and the processing of the FASTQ files, see supplementary methods. We used the HUMAnN2 pipeline^70^ with default settings for functional profiling of our samples, with the UniRef90 data base and ChocoPhlAn for alignment, and we renormalized our samples by library depth to copies per million. We used MetaCyc to obtain stratified and unstratified pathway abundances.

##### Statistical analysis of shotgun data

We calculated the predicted microbiota potency score for each sample and separately for neutrophils, lymphocytes and monocytes, by multiplying the abundances of taxa in each of the 124 samples with the corresponding posterior coefficients obtained from the PBSC inference. To distinguish the sets of metabolic functions that separate samples with positive and negative predicted potencies, we converted the pathway abundances into presence and absences profiles. We performed a linear discriminant analysis between positive and negative potency samples with a least squares solver and automatic shrinkage using the Ledoit-Wolf lemma using the sklearn package for the Python programming language^67^. To assess differences in the presence or absence of pathways between samples with positive and negative potency, we used Fisher’s exact test for each pathway.

## Supplementary Figures

**Figure S1:**
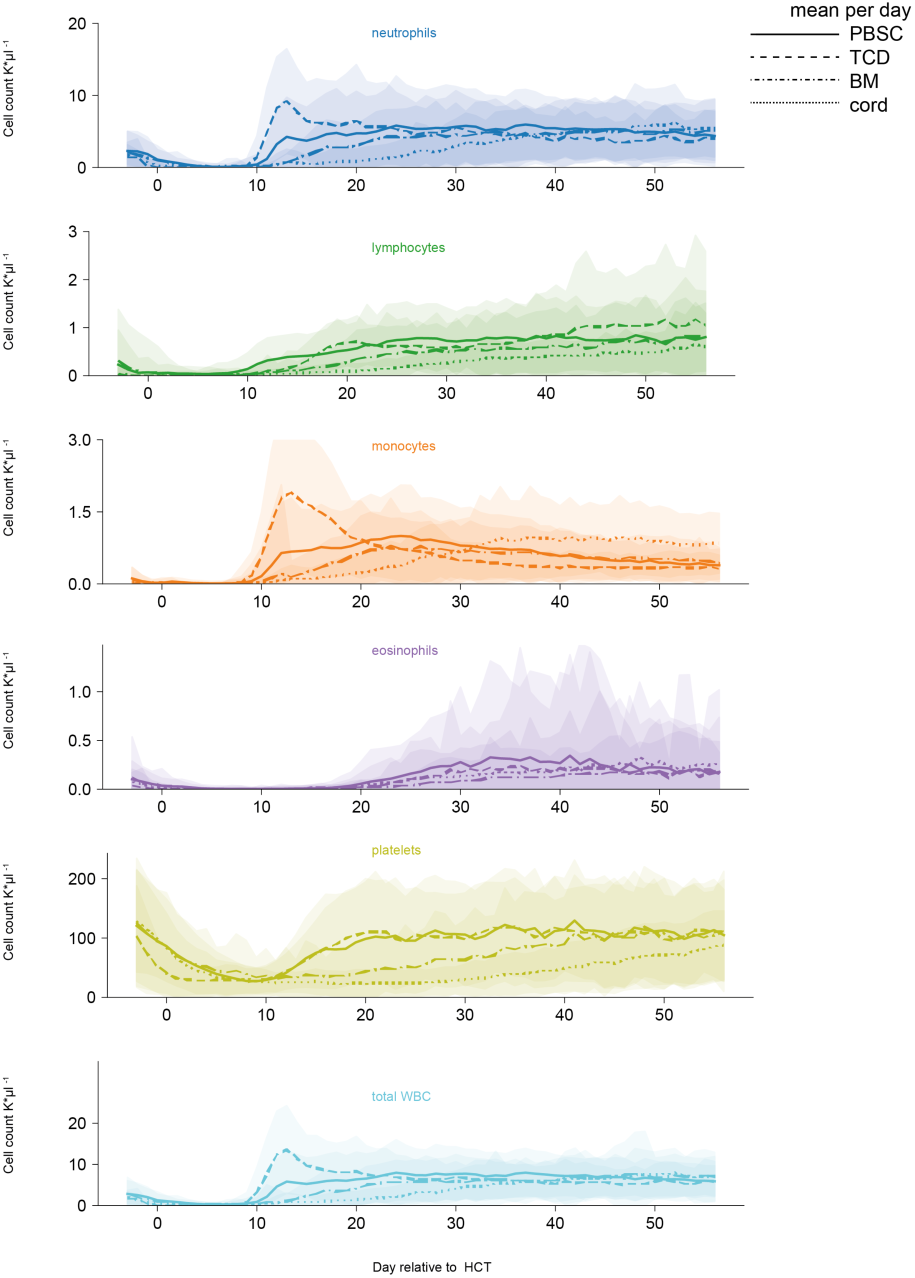
White blood cell counts and platelet counts per graft source over the first 100 days post HCT per day relative to HCT; lines: mean, shaded: ± standard deviations).

**Figure S2:**
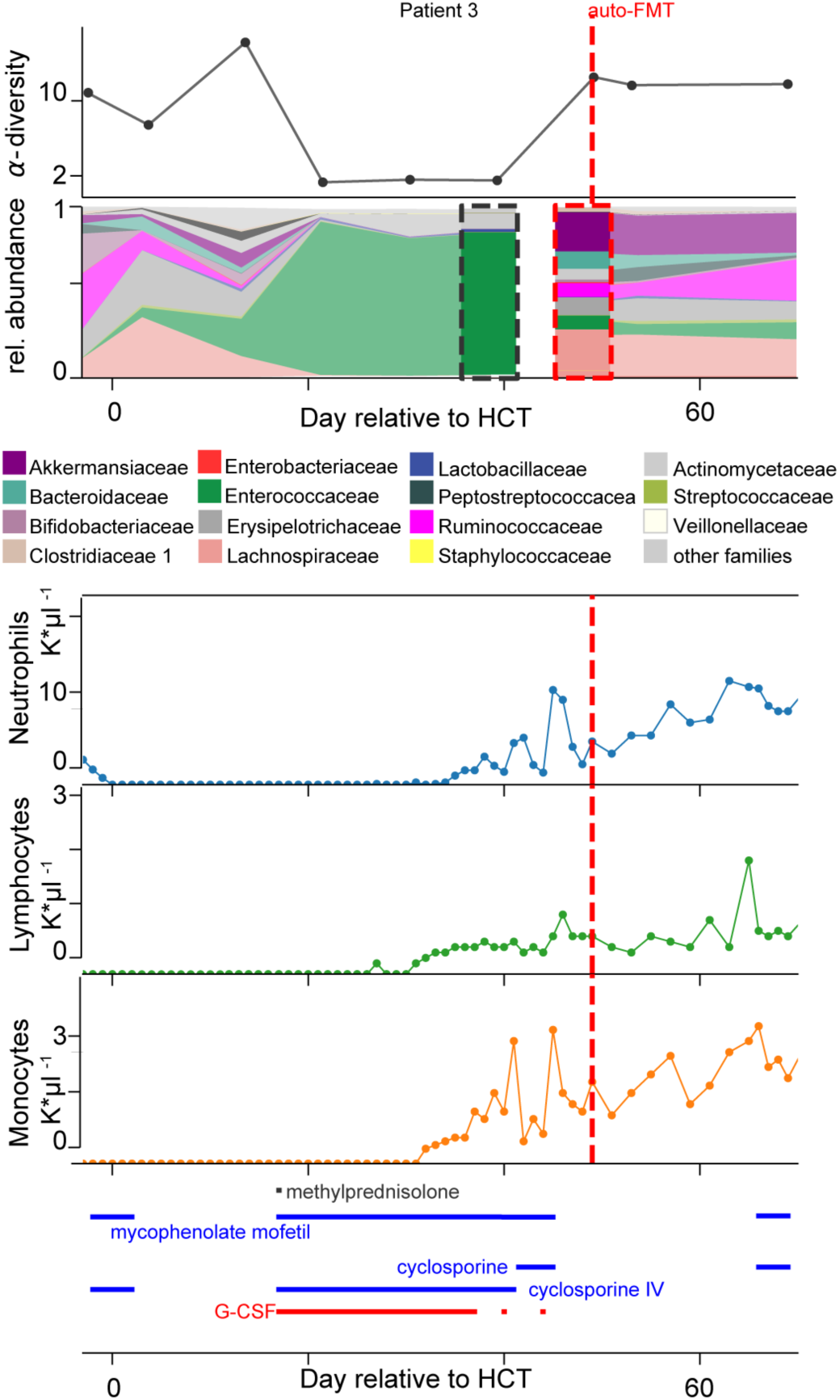
HCT patient who received an autologous fecal microbiota transplant (auto-FMT, dashed red line) that restored commensal microbial families and ecological diversity in the gut microbiota, with concurrent cell counts of peripheral neutrophils, lymphocytes and monocytes and immunomodulatory drug administrations.

**Figure S3:**
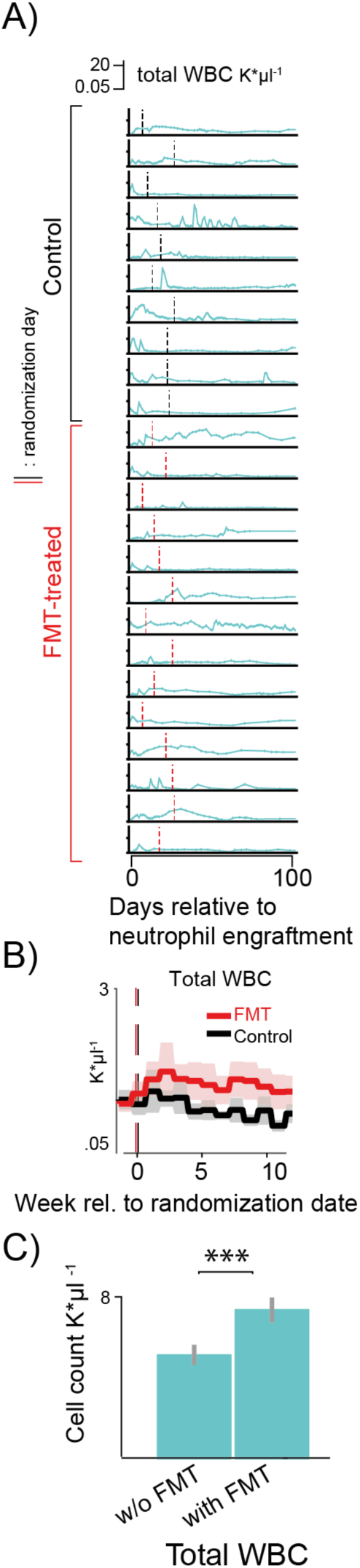
Total counts of white blood cells increased in FMT-treated patients in the weeks following the treatment compared to control patients. A) Total white blood cell counts in 24 patients enrolled in a randomized controlled trial to receive an autologous fecal microbiota transplant post-neutrophil engraftment (10 control: black vertical line, 14 FMT treated: red vertical line,). B) Weekly mean cell counts aligned to the date of randomization into FMT treatment arm (red) or control (black). Line: weekly mean, shaded region: 95%-CI C) Results from a linear mixed effects model with random effects per patient and per day relative to neutrophil engraftment confirms that the total white blood cell counts is higher in patients receiving auto-FMT after their treatment as compared to control patients after the randomization date, bars and confidence intervals for the averages of observed white blood cell counts without FMT and post-FMT (***: p<0.001).

**Figure S4:**
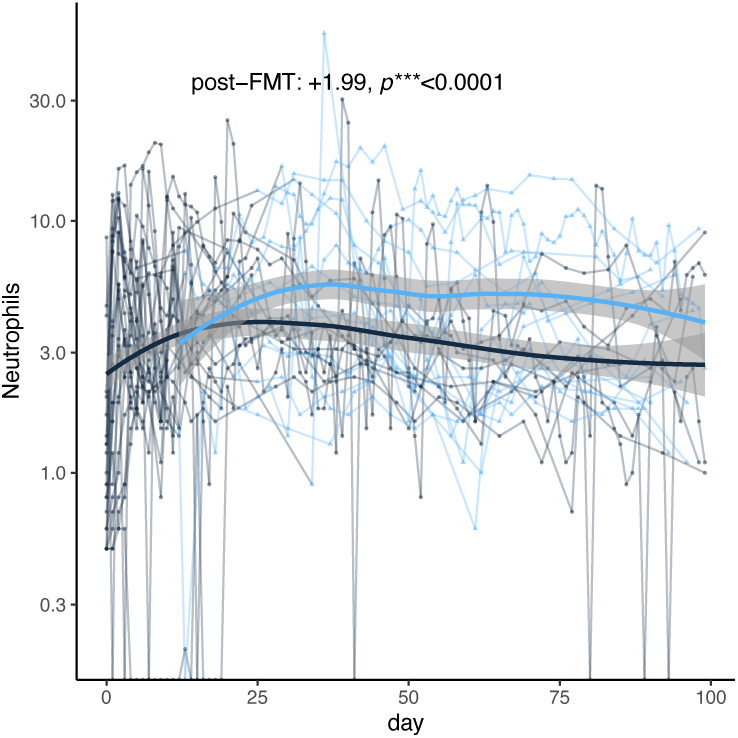
Neutrophil counts in 24. FMT trial patients. Thin lines: raw data (blue: post-FMT); thick black: mean per day, thick blue: mean+post-FMT coefficient. Means and confidence intervals (shaded region) from linear mixed effects model (methods).

**Figure S5:**
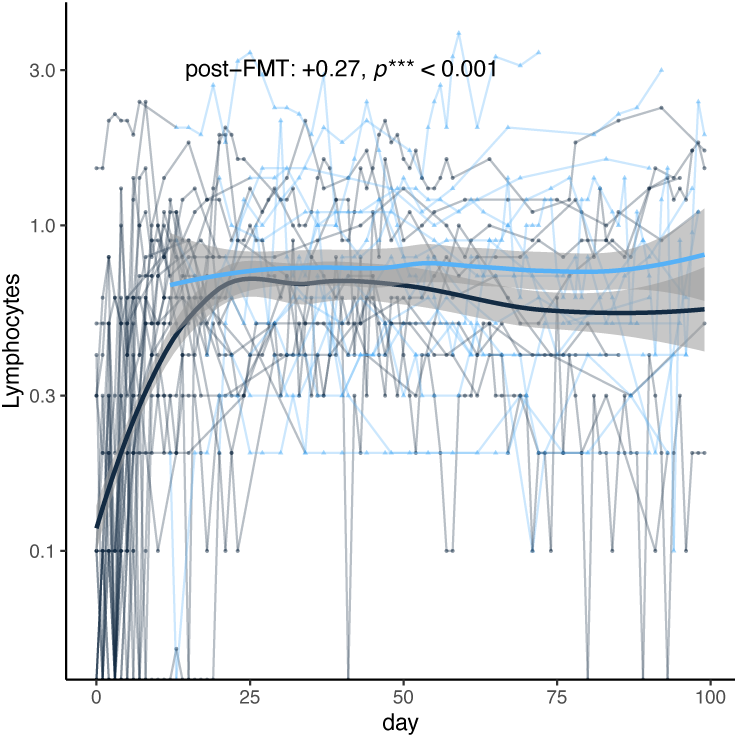
Lymphocyte counts in 24. FMT trial patients. Thin lines: raw data (blue: post-FMT); thick black: mean per day, thick blue: mean+post-FMT coefficient. Means and confidence intervals (shaded region) from linear mixed effects model (methods).

**Figure S6:**
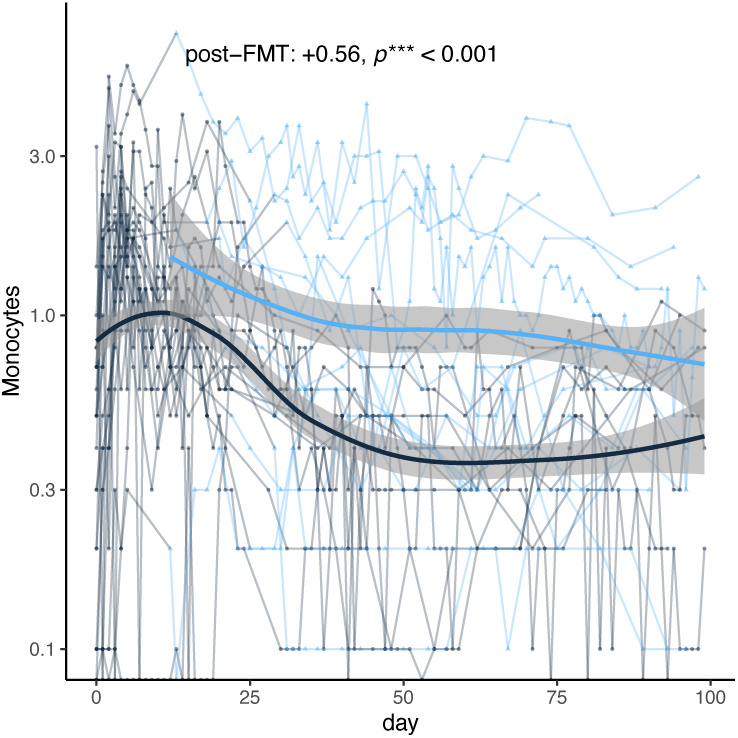
Monocyte counts in 24. FMT trial patients. Thin lines: raw data (blue: post-FMT); thick black: mean per day, thick blue: mean+post-FMT coefficient. Means and confidence intervals (shaded region) from linear mixed effects model (methods).

**Figure S7:**
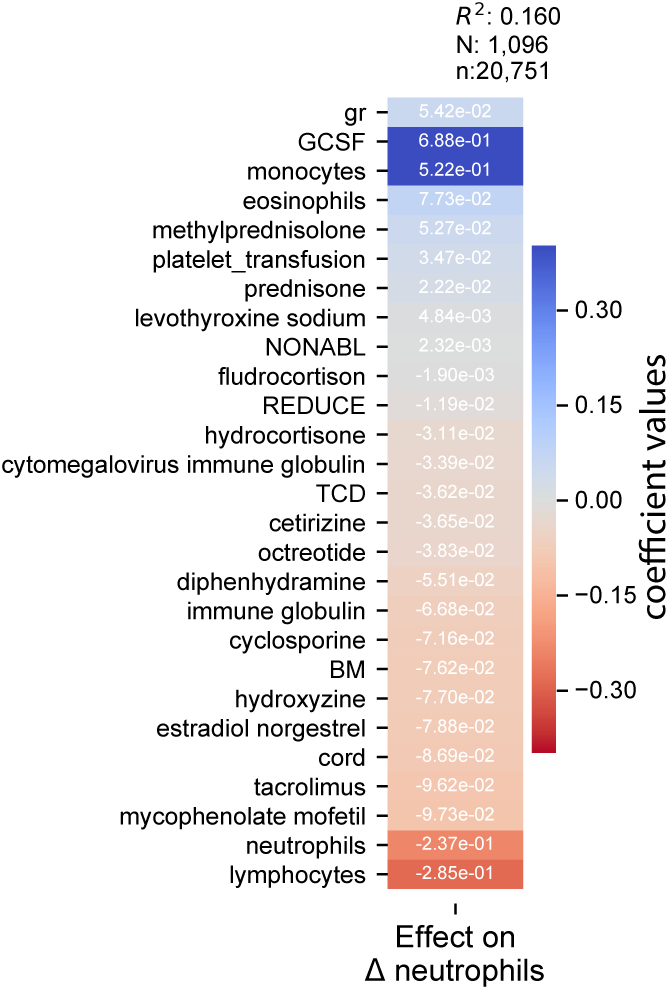
“Stage 1” regression on neutrophil dynamics on patients without microbiome data. Coefficients from 10-fold cross-validated elastic net regression daily changes in neutrophils. gr: intercept; TCD: T-cell depleted graft (ex-vivo) by CD34+selection; PBSC: peripheral blood stem cells; BM: bone marrow; cord: umbilical cord blood; NONABL: Nonmyeloablative; REDUCE: reduced-intensity conditioning regimen; F: female; N: patients, n: samples (daily changes in neutrophils).

**Figure S8:**
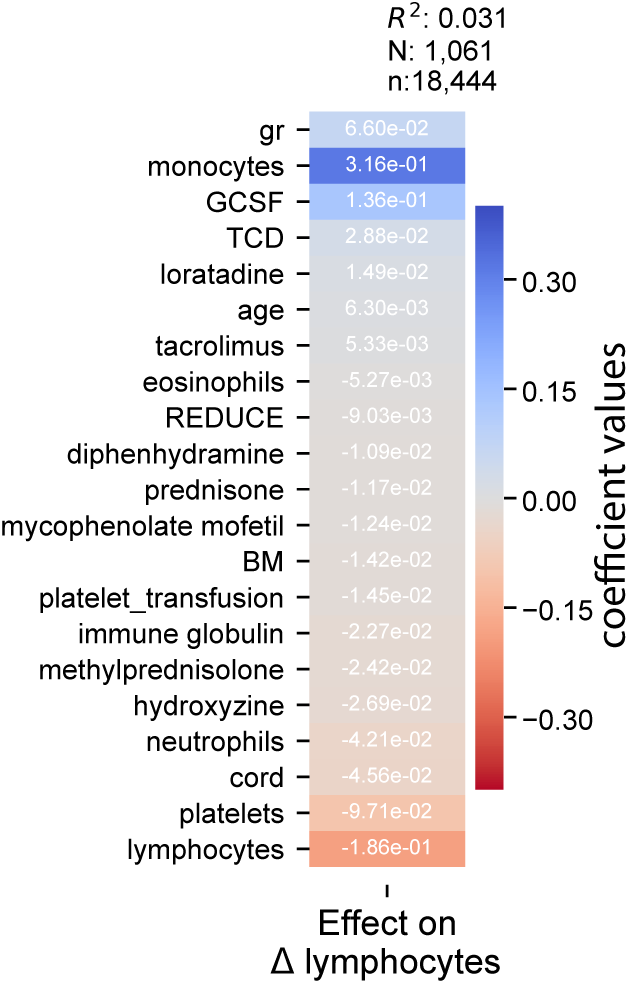
“Stage 1” regression on lymphocytes dynamics on patients without microbiome data. Coefficients from 10-fold cross-validated elastic net regression daily changes in lymphocytes. gr: intercept; TCD: T-cell depleted graft (ex-vivo) by CD34+selection; PBSC: peripheral blood stem cells; BM: bone marrow; cord: umbilical cord blood REDUCE: reduced-intensity conditioning regimen; F: female. N: patients, n: samples (daily changes in lymphocytes).

**Figure S9:**
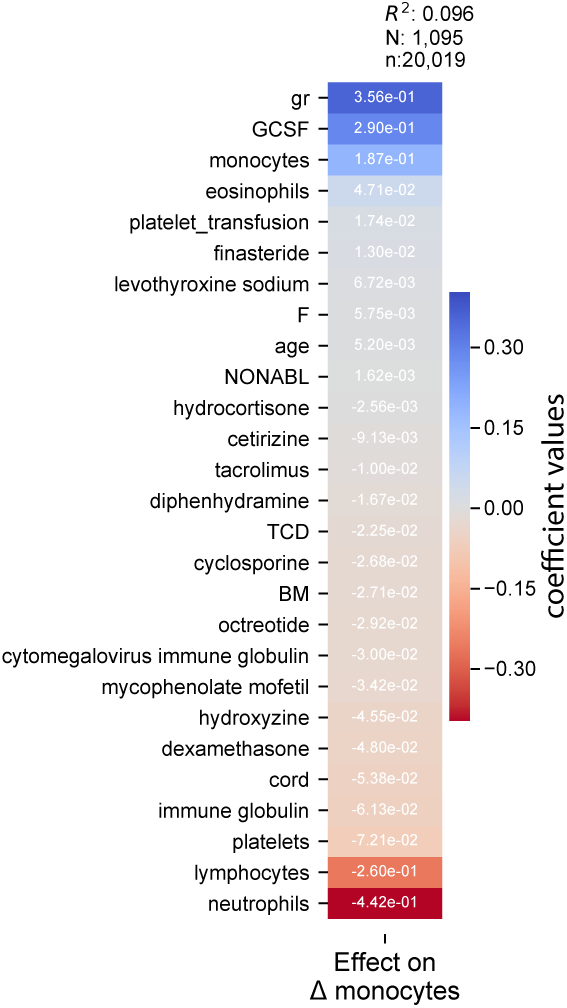
“Stage 1” regression on lymphocytes dynamics on patients without microbiome data. Coefficients from 10-fold cross-validated elastic net regression daily changes in lymphocytes. gr: intercept; TCD: T-cell depleted graft (ex-vivo) by CD34+selection; PBSC: peripheral blood stem cells; BM: bone marrow; cord: umbilical cord blood REDUCE: reduced-intensity conditioning regimen; F: female. N: patients, n: samples (daily changes in lymphocytes).

**Figure S10:**
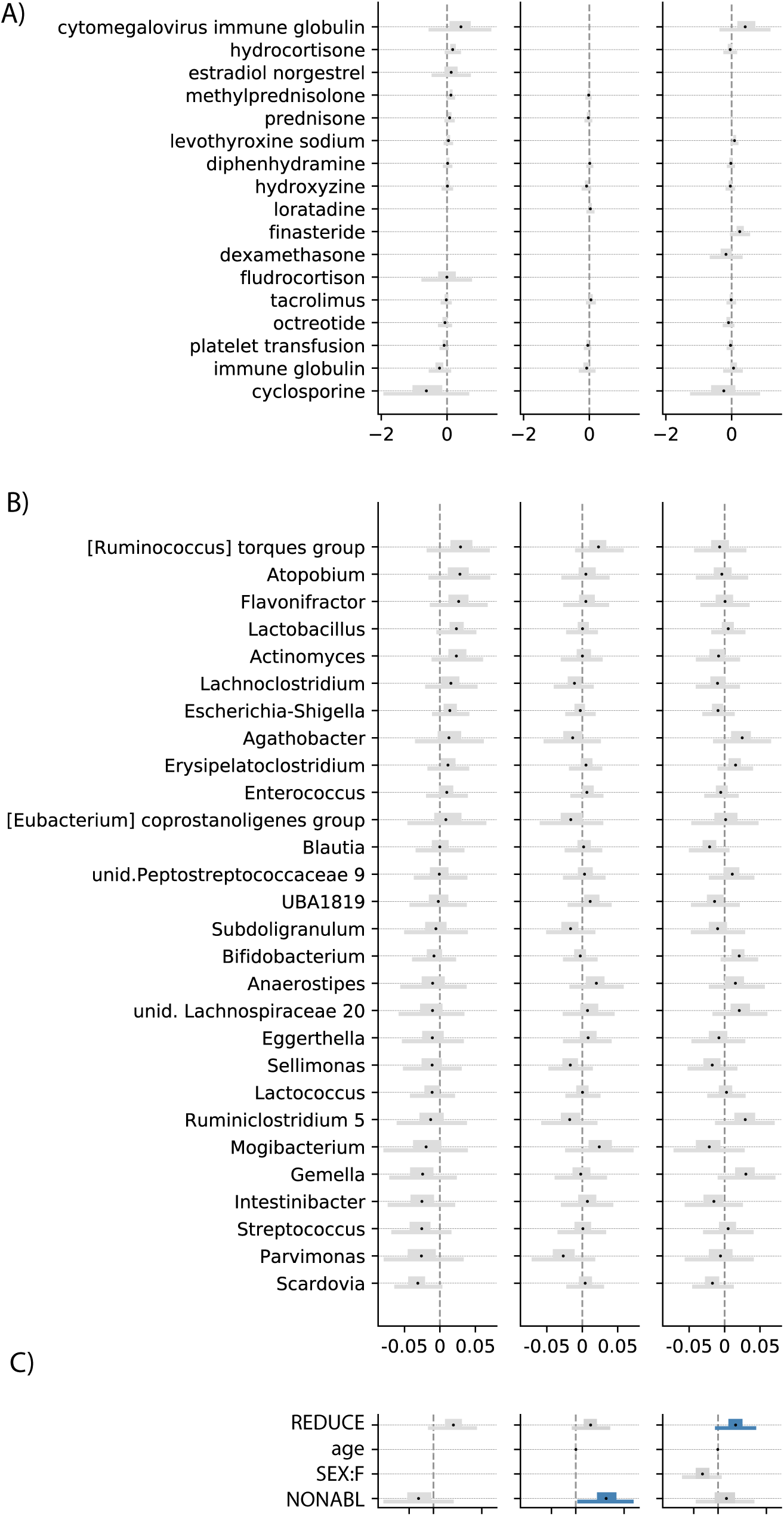
Additional coefficient estimates of medications (A), additional genera (B) and metadata (C) from the Bayesian regression, see also Figure 3. REDUCE: reduced-intensity conditioning regimen; NONABL: non-myeloablative conditioning regimen. F: female

**Figure S11:**
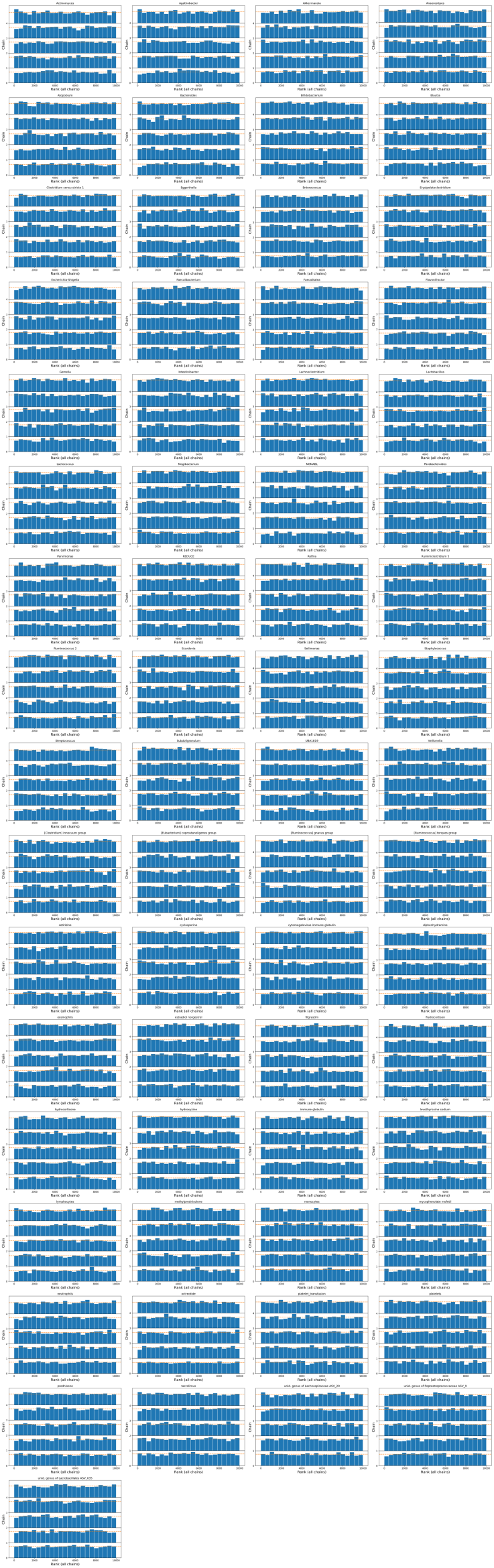
Posterior sampling converged. Histograms of the ranked posterior draws from the model of neutrophil dynamics in PBSC patients (ranked over all chains), plotted separately for each chain (see supplementary methods), show no substantial differences between chains^71^.

**Figure S12:**
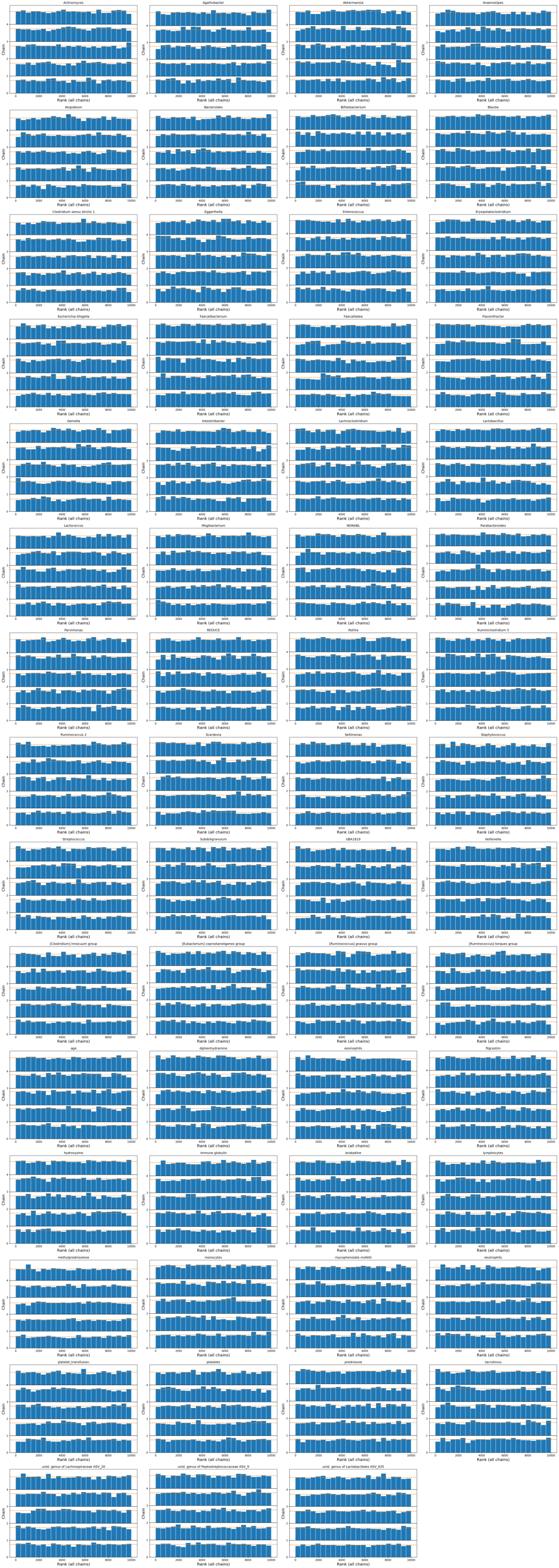
Posterior sampling converged. Histograms of the ranked posterior draws from the model of lymphocyte dynamics in PBSC patients (ranked over all chains), plotted separately for each chain (see supplementary methods), show no substantial differences between chains^71^.

**Figure S13:**
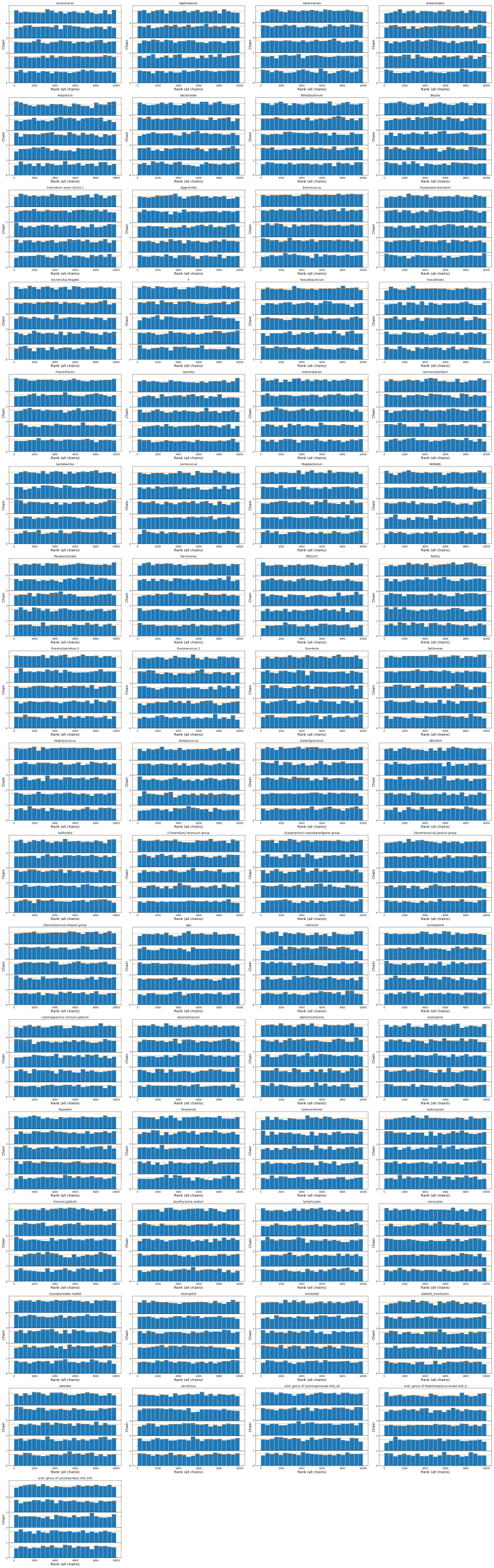
Posterior sampling converged. Histograms of the ranked posterior draws from the model of monocyte dynamics in PBSC patients (ranked over all chains), plotted separately for each chain (see supplementary methods), show no substantial differences between chains^71^.

**Figure S14:**
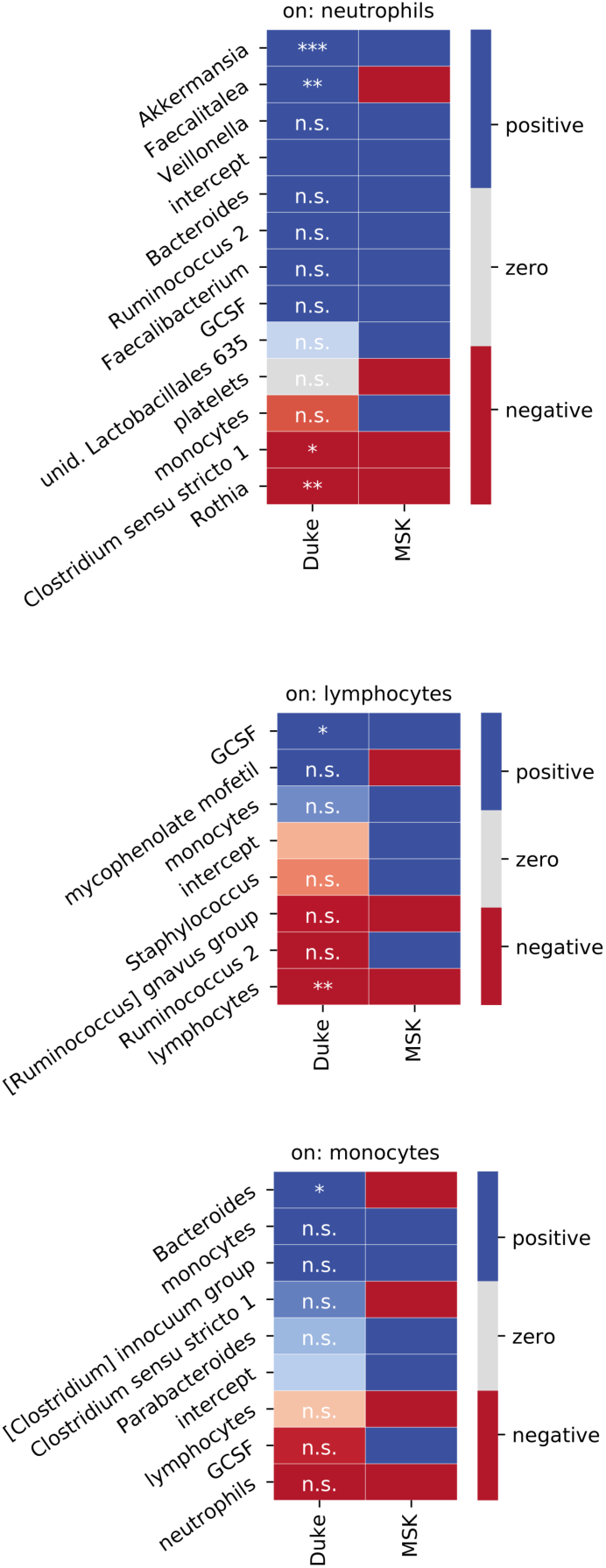
Validation analysis of predictors on white blood cell dynamics using data from patients treated at DukeHealth. Individual univariate regressions of microbiome and clinical predictors identified in stage 2 of our analysis on daily changes in neutrophils, lymphocytes and monocyte. Bonferroni corrected p-values: ***<0.001, **<0.01, *<0.05; p>0.05: n.s. Sign of coefficients from MSK PBSC patients for comparison.

**Figure S15:**
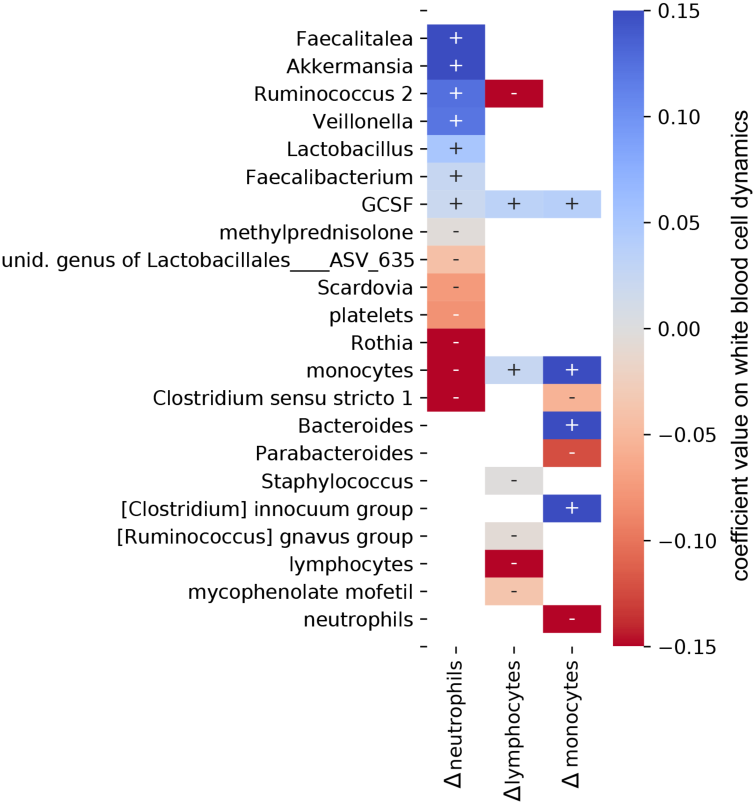
Validation analysis of predictors on white blood cell dynamics using data from patients treated at DukeHealth. Partial least squares regression of microbiome and clinical predictors identified in stage 2 of our analysis on daily changes in neutrophils, lymphocytes and monocyte.

**Figure S16:**
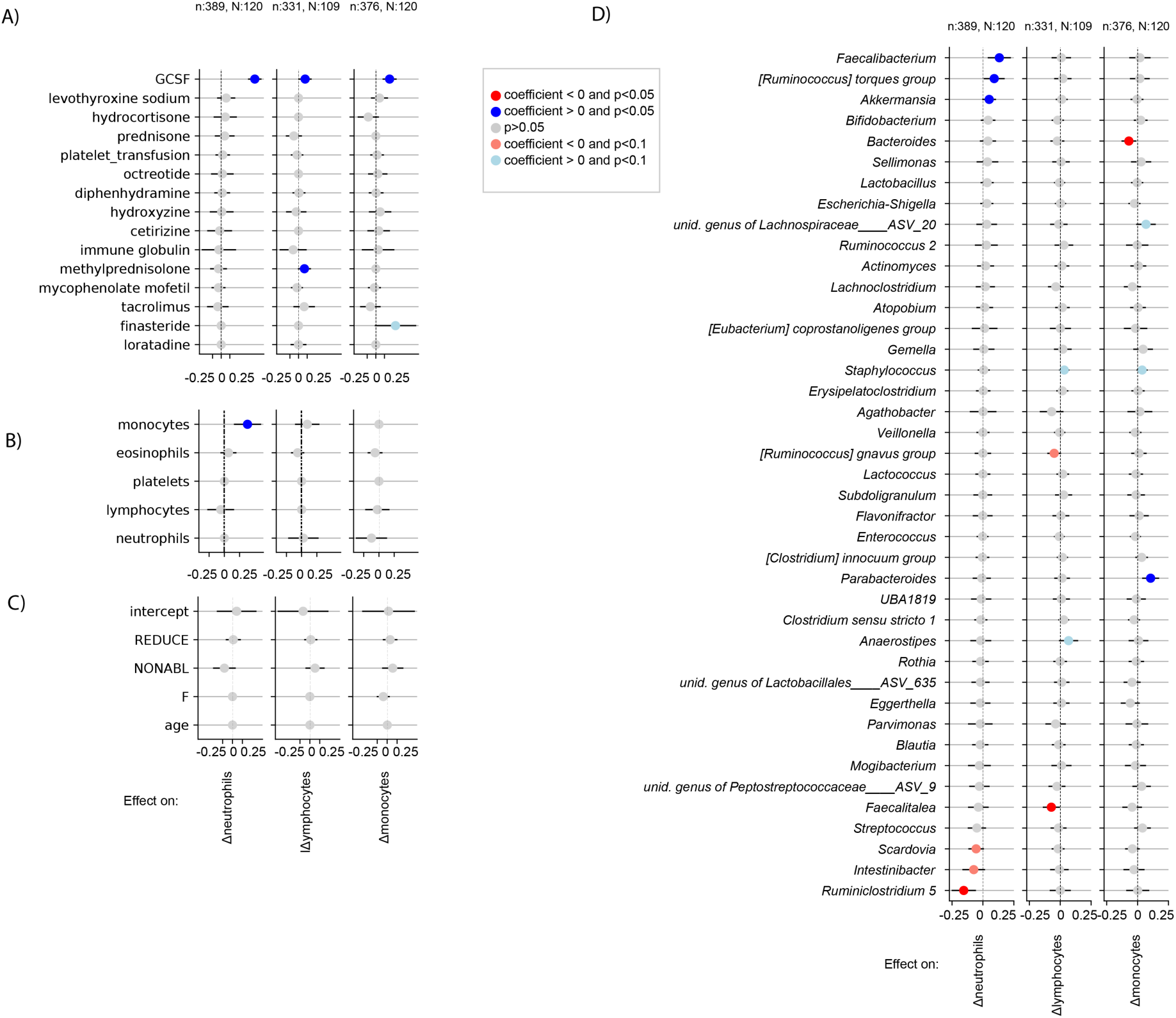
Validation analysis of the main model using absolute bacterial abundances as predictors instead of relative abundances in Figure 3. Results show coefficients from a least squares regression for medications (A), white blood cell feedbacks (B) metadata (C) and total genus abundances (D) of neutrophil, lymphocyte and monocyte daily log-changes. This was only possible for only a subset of the data for which we obtained absolute bacterial abundance estimates (methods), n: samples, N: patients.

**Figure S17:**
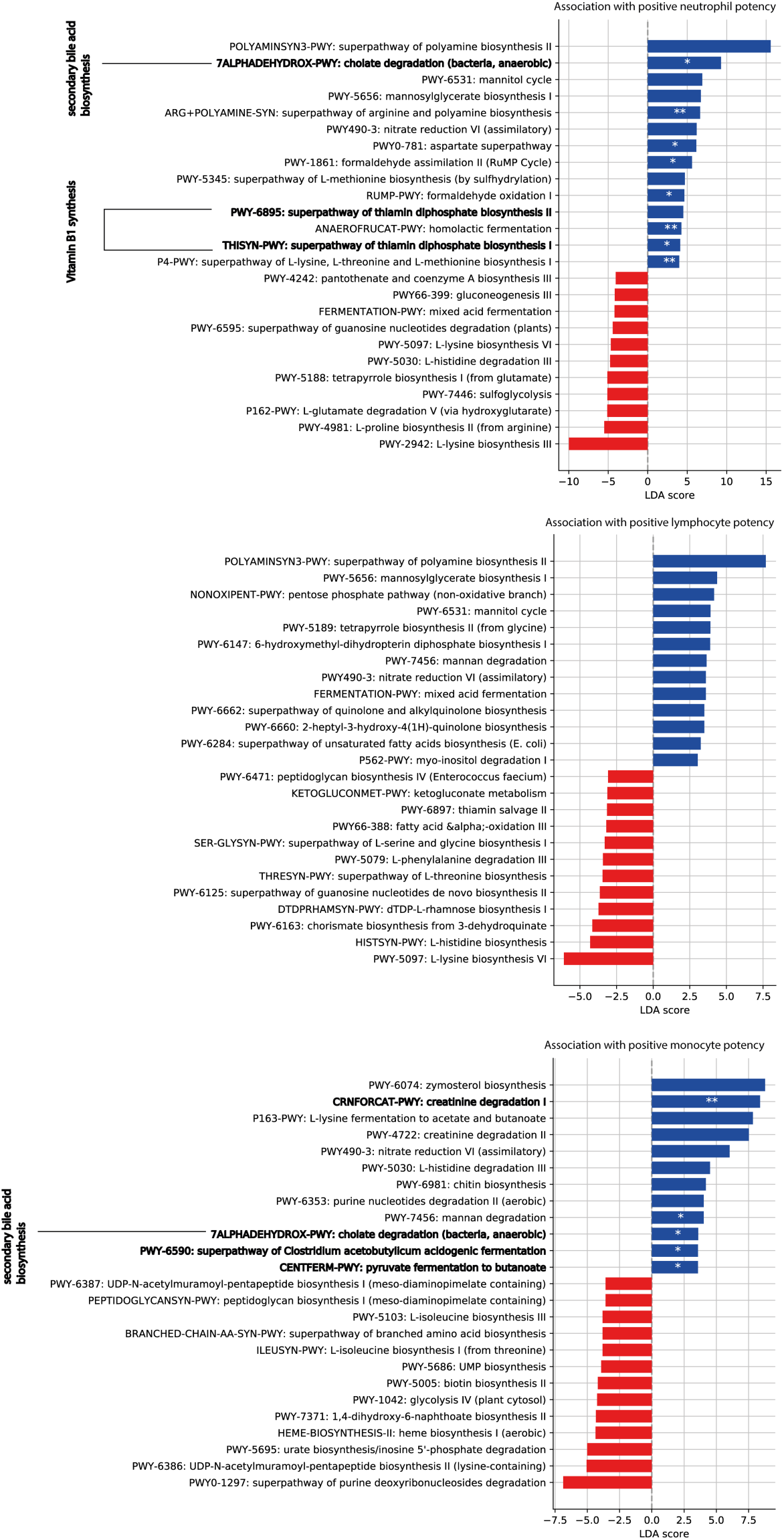
Functional analysis of microbiota samples. To distinguish samples predicted to increase rates of white blood cells, a microbiota potency score was calculated from posterior coefficients (Figure 3, methods) and the relative abundance of taxa in samples. Bars show linear discriminant analysis (LDA) scores of MetaCyc pathway profiles from 124 shotgun sequenced samples that distinguished positive and negative potency samples the most (LDA-score magnitude in the 95^th^ percentile). Highlighted pathways are discussed in the main text. For each pathway, we tested differences between positive and negative potency samples using Fisher’s exact test; p-value <0.001: ***, <0.01:**, <0.05:*.

**Figure S18:**
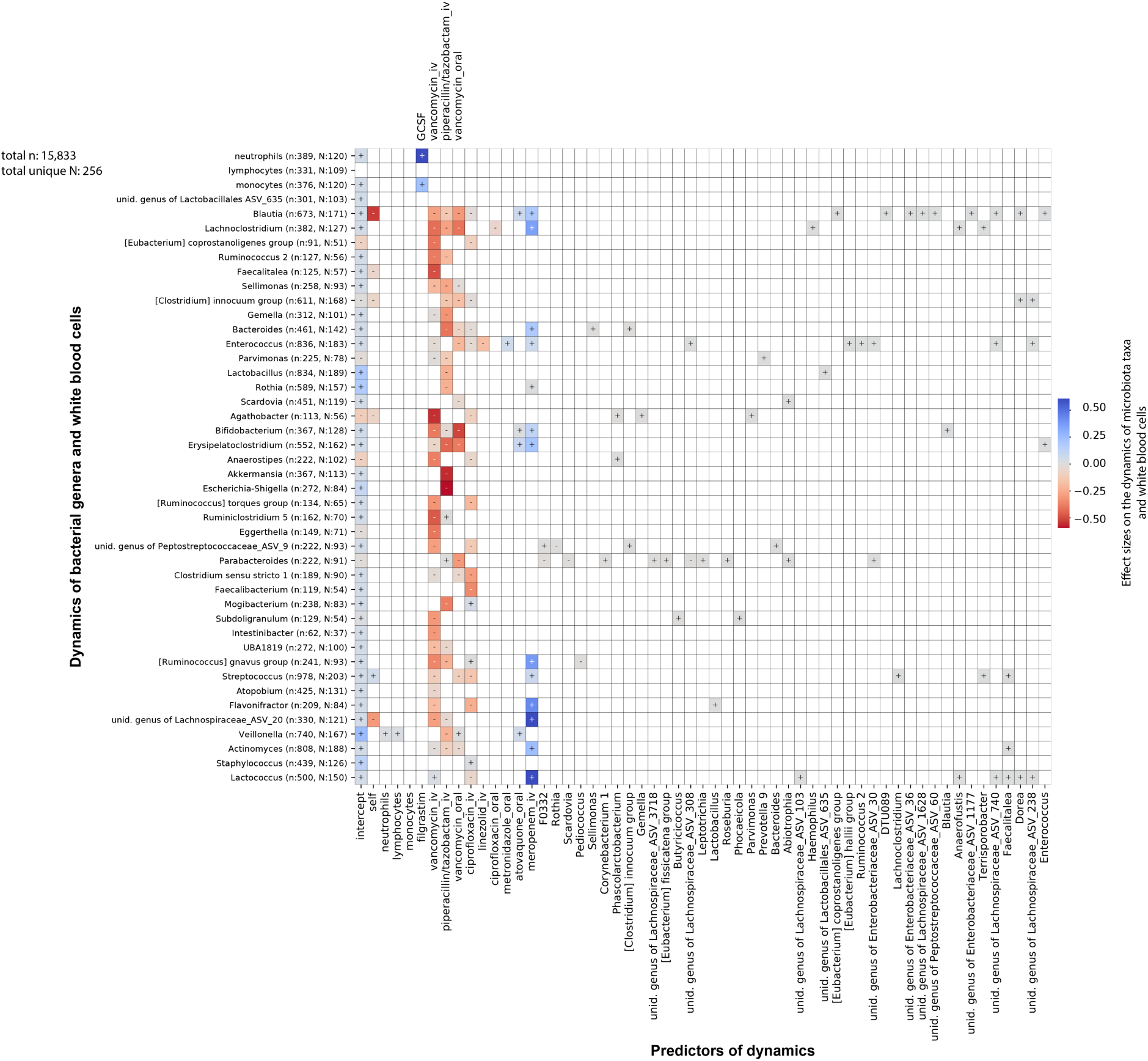
Jointly inferred association network between white blood cell and bacterial genus dynamics (methods). Strong regularization yields few non-zero coefficients and antibiotics dominate the dynamics.

**Figure S19:**
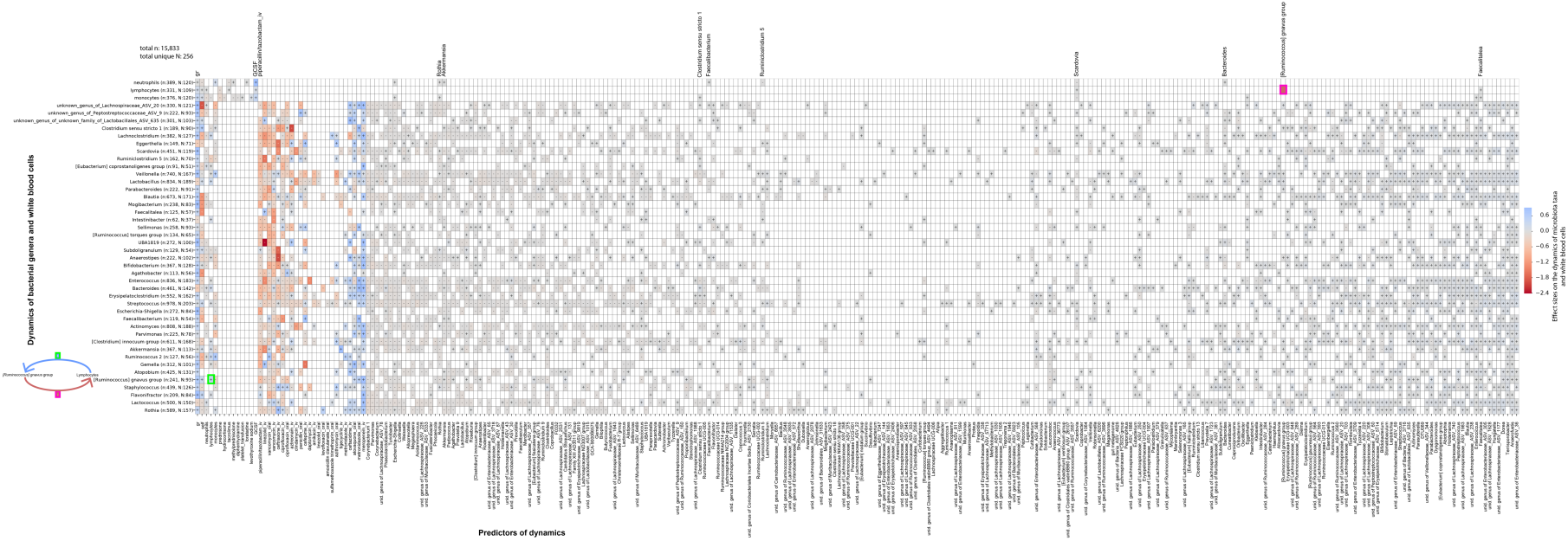
Jointly inferred association network between white blood cell and bacterial genus dynamics with reduced regularization (methods) indicates potential bidirectional feedbacks, e.g. between lymphocytes and *[Ruminococcus] gnavus group* (highlighter green boxes, and cartoon).

**Figure S20:**
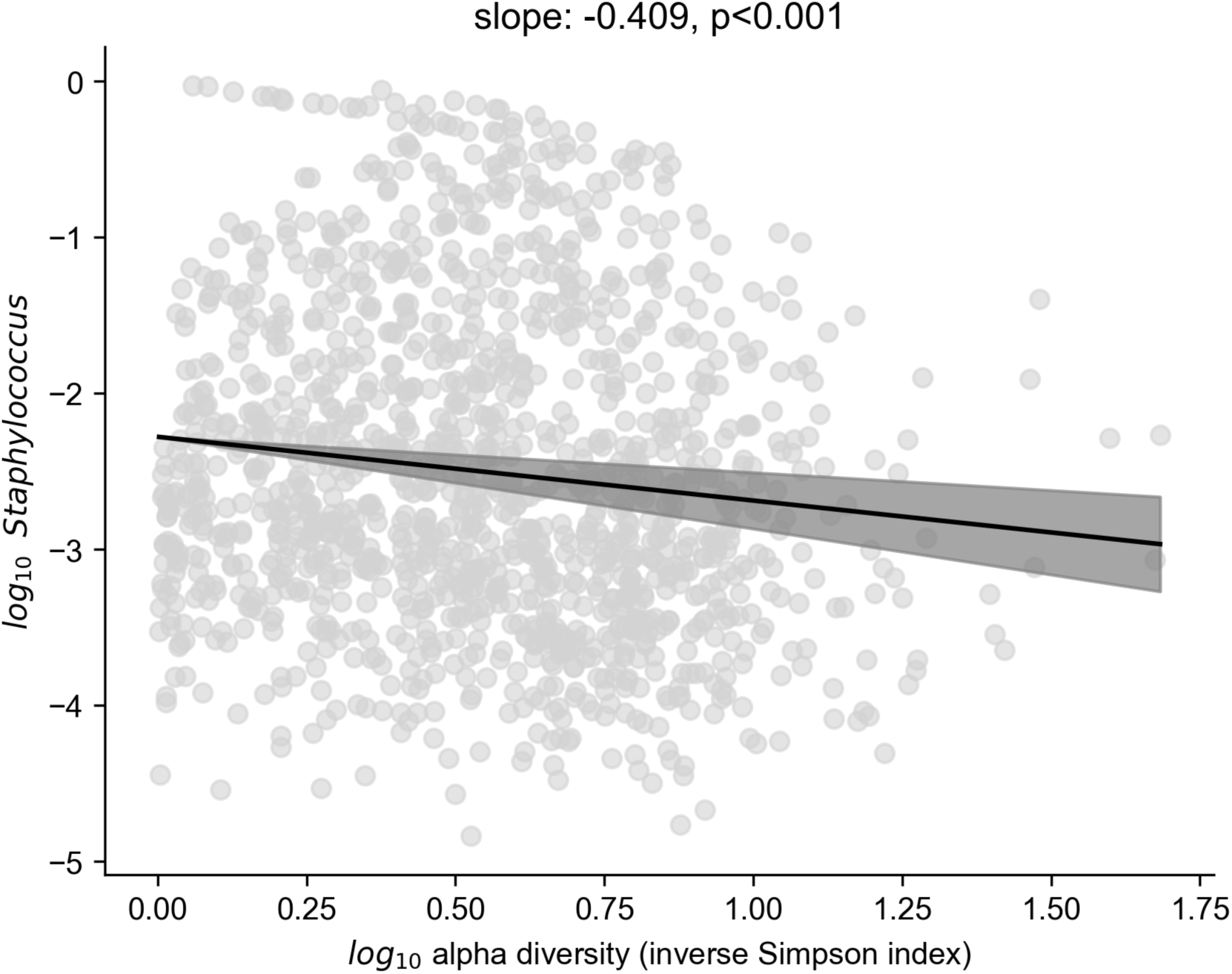
The relative non-zero abundance of Staphylococcus is inversely related to microbiome alpha diversity, shaded: 95% confidence intervals.

**Figure S21:**
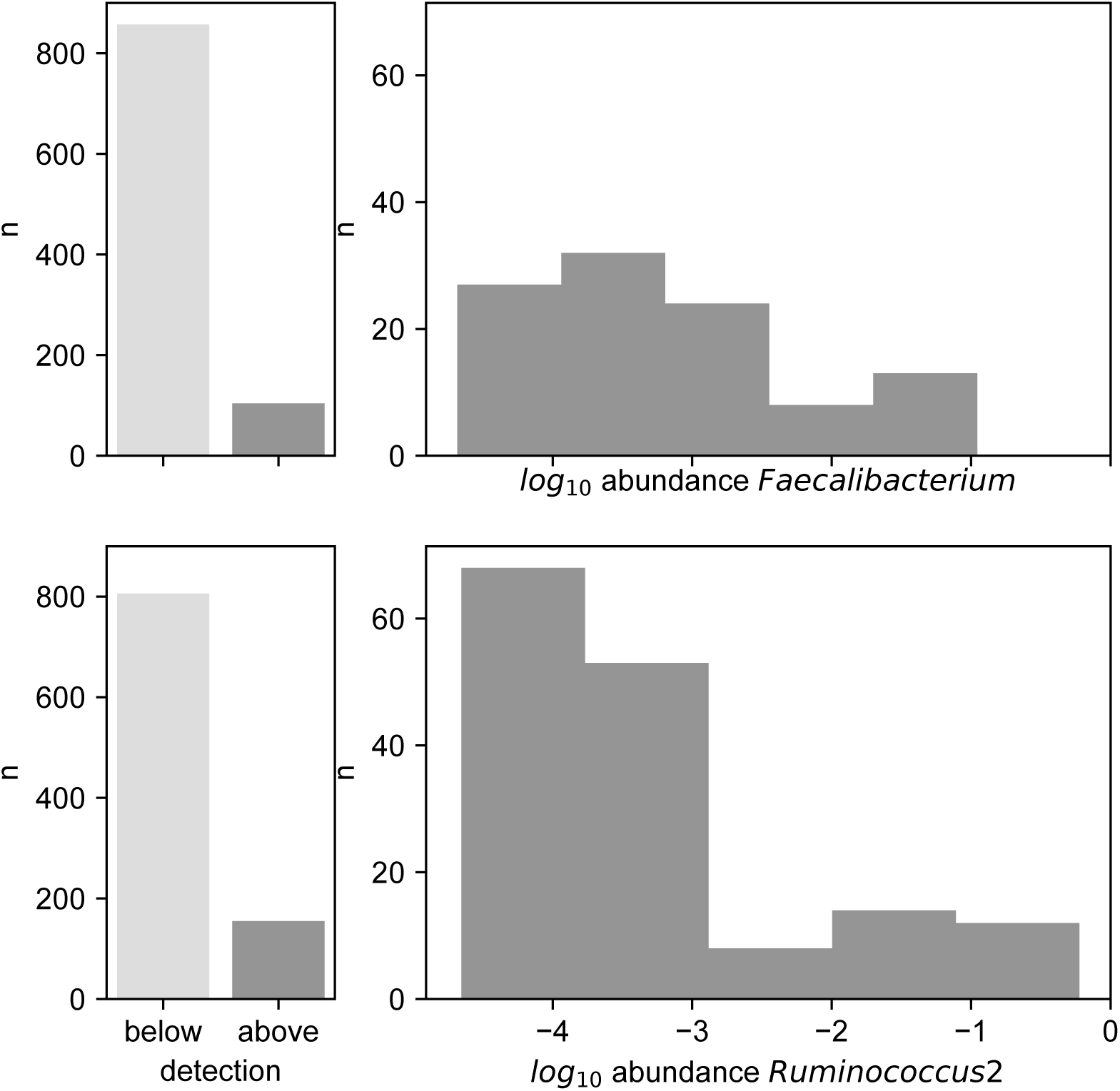
Abundance profiles of the two genera, *Faecalibacterium* and *Ruminococcus 2*, most strongly associated with white blood cell increase; number of times detected (left) and *log*_*10*_ abundance distribution when above detection (right).

**Figure S22:**
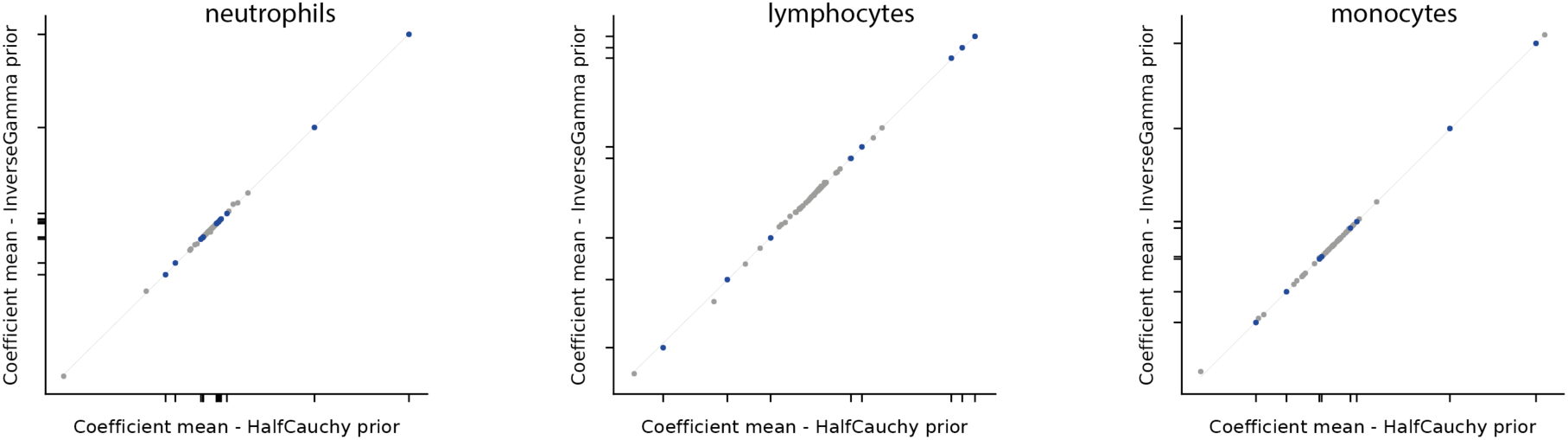
Posterior association coefficients do not depend on the choice of prior for σ in the main Bayesian model. Plotted are the posterior means from our main analysis against the equivalent inference with an inverse Gamma prior (alpha=1, beta=1).

**Table S1:** Data set summary and patient characteristics. HCT-graft types: TCD: T-cell depleted graft (*ex-vivo*) by CD34+selection; PBSC: peripheral blood stem cells; BM: bone marrow; cord: umbilical cord blood; Conditioning intensity: Bacigalupo classification, graded categories from most to least intense (ABLATIVE, REDUCE, NONABL).

|  |  |  |
| --- | --- | --- |
| **patients** |  | 2,926 |
| **HCT therapies\*** |  | 3,060 |
| **blood samples** | total | 450,635 |
|  | between HCT-day -21 and HCT-day 183 | 193,396 |
| **Disease** | Leukemia | 1,635 |
|  | Non-Hodgkin's Lymphoma | 415 |
|  | Multiple Myeloma | 170 |
|  | Hodgkin's disease | 88 |
|  | other | 752 |
| **HCT graft type** | TCD | 1,106 |
|  | PBSC unmodified | 959 |
|  | BM unmodified | 617 |
|  | cord | 378 |
| **Conditioning intensity** | ABLATIVE | 65% |
|  | REDUCE | 21% |
|  | NONABL | 13% |
| **Gender** | M | 58% |
|  | F | 42% |
| **Age of adults (years)** | 25%-tile | 39 |
|  | mean | 50 |
|  | 75%-tile | 62 |
| **Microbiome samples** | total | 12,633 |
|  | - from patients with blood data | 10,680 |
|  | - of those, post engraftment | 4,179 |
|  | - of those with daily change in WBC | 2,615 |
|  | patients with microbiome sample | 1,290 |

\*) some patient received several HCTs

**Table S2:** Patient and HCT characteristics of 24 patients enrolled in the randomized controlled FMT trial.

|  | **control** | **FMT treated** |
| --- | --- | --- |
| **N patients** | 10 | 14 |
| **ABLATIVE** | 6 | 7 |
| **REDUCE** | 4 | 7 |
| **BM unmodified** | 1 | 3 |
| **PBSC unmodified** | 3 | 4 |
| **TCD** | 5 | 3 |
| **cord** | 1 | 4 |

**Table S3:**
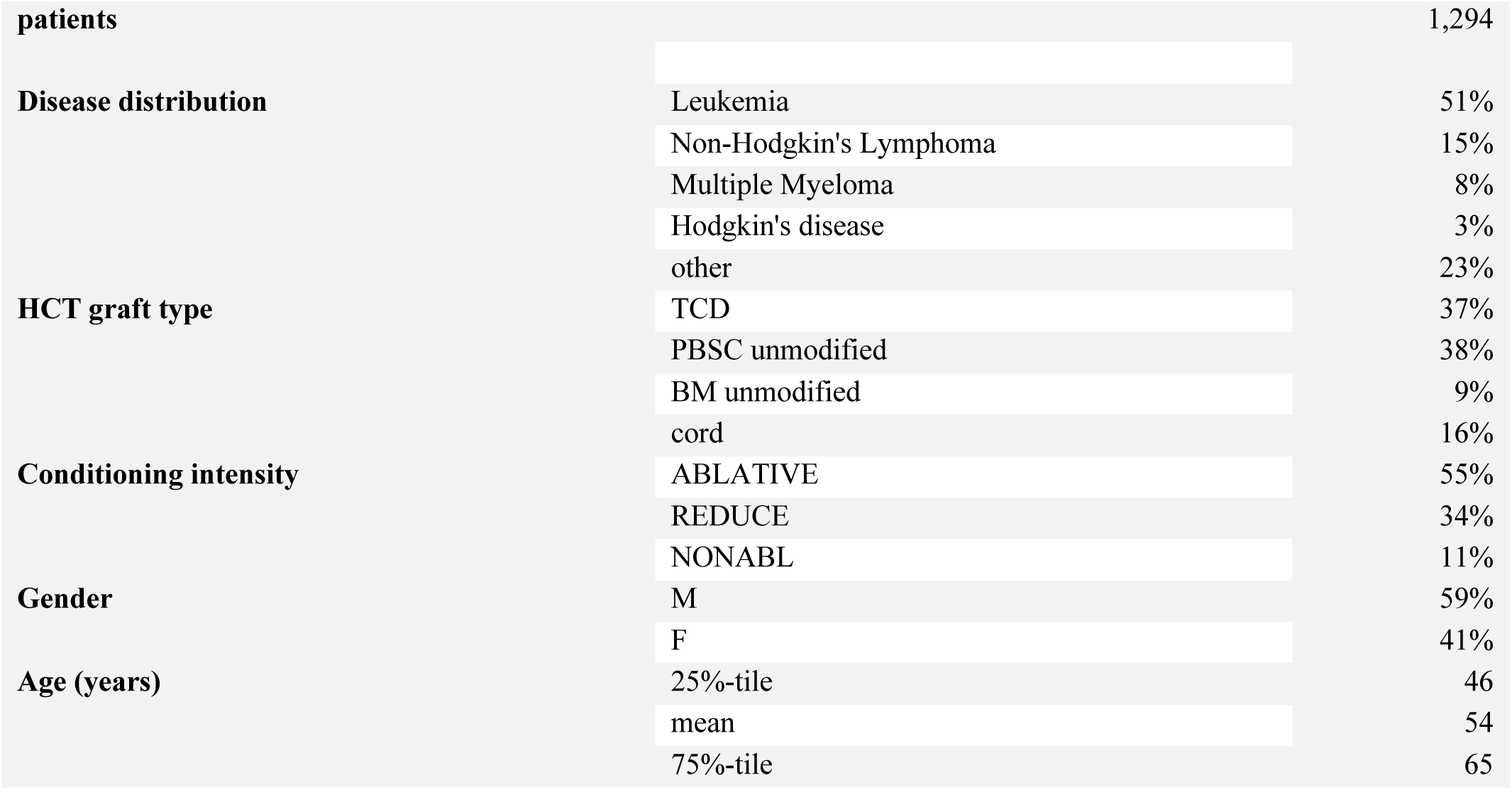
Patient and HCT characteristics of the subset of patients who donated microbiota samples.

**Table S4:**
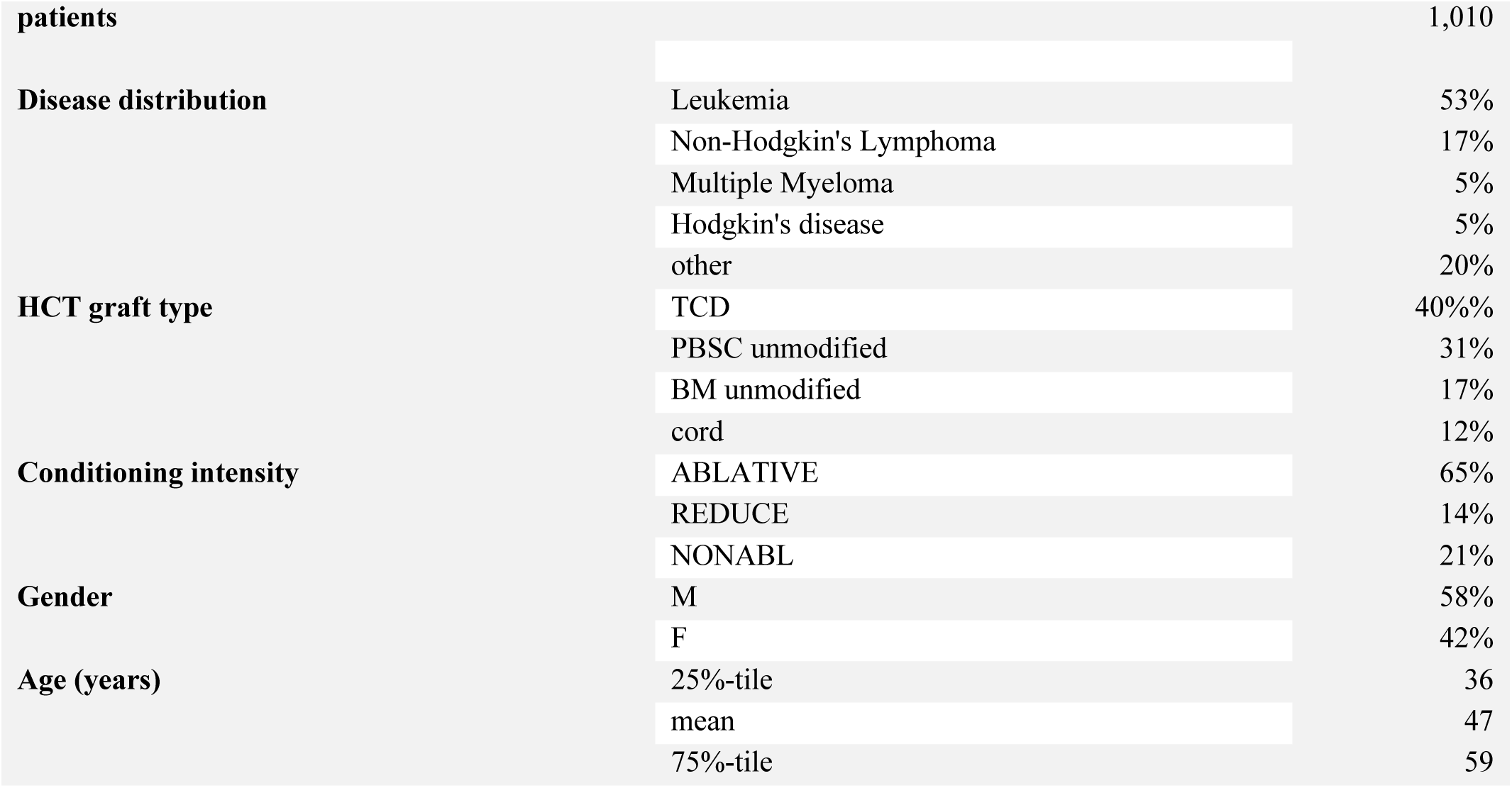
Patient and HCT characteristics of the subset of patients who did not donate microbiota samples.

**Table S5:**
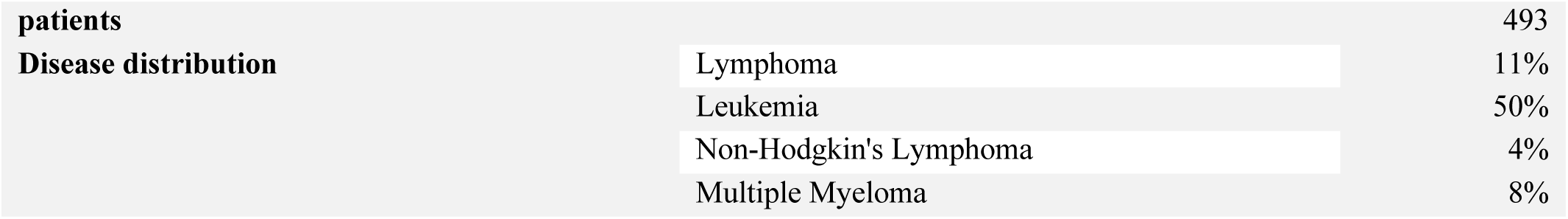
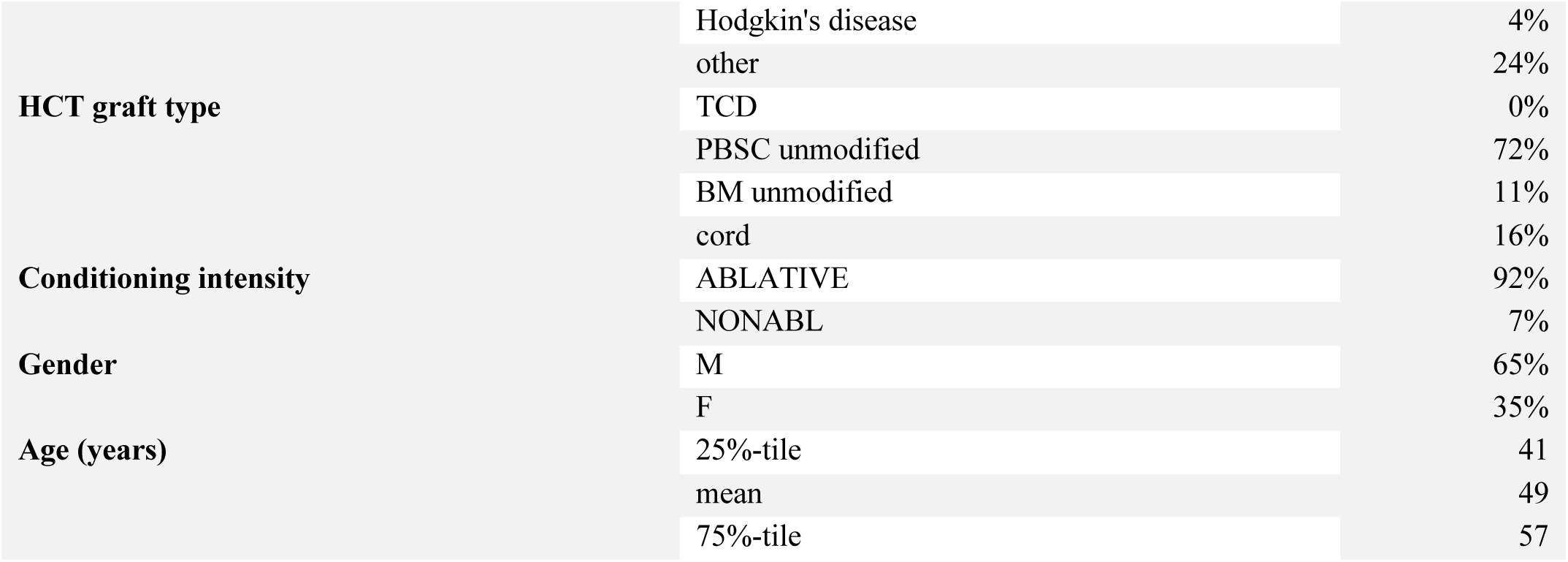
Patient and HCT characteristics of the Duke University patient cohort.

## Data availability

The data used in our study is organized in supplementary tables (data-tables.zip), with corresponding filenames (italic):

1. *cGENUS*.*csv*: relative taxon abundances in fecal microbiota samples from 12,633 stool samples
2. *cHCTMETA*.*csv*: HCT characteristics
3. *cINFECTIONS*.*csv*: positive blood culture results
4. *cMISAMPLES*.*csv*: NCBI SRA accession number, diversity (inverse Simpson index), total 16S (where available), stool consistency for each fecal microbiota sample
5. *cMED*.*csv*: medication data
6. *cPIDMETA*.*csv*: anonymized patient demographics
7. *cWBC*.*csv*: absolute counts of neutrophils, lymphocytes, monocytes, eosinophils, and platelets with indication if included in analyses
8. *cDUKE GENUS*.*csv*: relative taxon abundances in fecal microbiota samples from 12,633 stool samples
9. *cDUKE WBC*.*csv*: absolute counts of neutrophils, lymphocytes, monocytes, eosinophils, and platelets with indication if included in analyses
10. *cDUKE MED*.*csv*: medication data
11. *cFMT_analysis*.csv: convenience table for Figure 2

## Code availability

The relevant scripts for stage 1, stage 2, and the model assessing the effect of FMT on white blood cell counts are on github:https://github.com/jsevo/wbcdynamics_microbiome.

