## Supplementary methods for "An association between the gut microbiota and immune cell dynamics in humans"

**FMT procedure**

Patient sample collection protocols were approved by the Memorial Sloan Kettering Cancer Center Institutional Review and Privacy Board (ClinicalTrials.gov identifier: NCT02269150) and described in full in the original publication. Briefly, patients’ stool was collected when they first entered the clinic. We chose autologous, as opposed to feces from a heterologous donor, because of potential safety concerns. The stool was tested for the presence of potential intestinal pathogens including *C. difficile* and frozen (−80°C). If a patient was randomized to receive treatment after engraftment of neutrophils, the thawed sample was re-administered via an enema. Due to the strenuous nature of this procedure, it was deemed unethical to administer a mock enema to control patients. Subjects whose pre–allo-HSCT feces demonstrated low microbial diversity (IS index < 2.0) or tested positive for the presence of an intestinal pathogen, for example, *C. difficile*, were excluded from randomization. After successful hematopoietic cell engraftment (three consecutive blood neutrophil counts ≥500 per mm3), subjects underwent testing of a fecal specimen collected after engraftment to determine the presence of the Bacteroidetes phylum via quantitative PCR. Individuals with low abundance of Bacteroidetes (<0.1% total 16S) were eligible to proceed to randomization and treatment. Eligible subjects were 1:1 randomized to undergo auto-FMT with the subject’s stored pre–allo-HSCT feces versus no fecal transplantation. Randomization was stratified by cord blood source versus non–cord blood source. Subjects could be randomized within a 28-day window after engraftment. Subjects who were critically ill or required prolonged microbiota-perturbing antibiotics through the designated 28-day period were excluded from randomization.

**Alternative analysis of FMT effect on white blood cell counts**

In the main text we considered a random effect per day post neutrophil engraftment. Our choice of modeling day_i as a random intercept term was to allow the cell counts to follow any function of day post FMT. Alternatively, we conducted another analysis now with day as a linear predictor:$y_{ij}= \beta_{0}+day+ {armpost*FMT}_{ij}+{patient}_{j}+\varepsilon_{ij}, i=0,\ldots,D, j=1,\ldots,P$,

This reduced significance but did not alter our main conclusions: FMT associated with increased neutrophils (coefficient FMT: 1.44, p=0.002), lymphocyte (coefficient FMT: 0.08, p=0.13), and monocytes (coefficient FMT: 0.34, p<0.001).

**Survival analysis**

We sought to analyze 3-year survival following neutrophil engraftment in the patient cohort analyzed in our main analyses. We observed 668 deaths during that period (see Kaplan Meier plot).

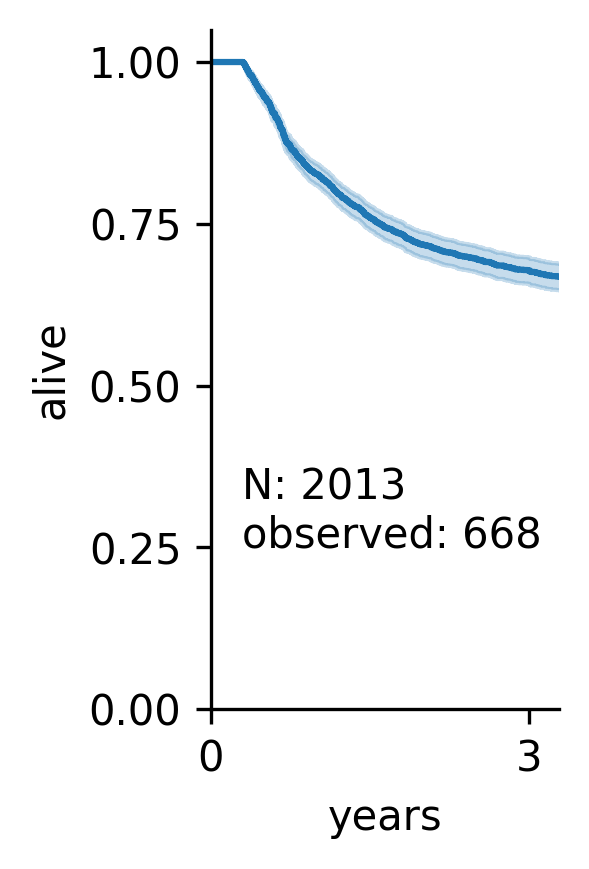

We used a Cox proportional hazards model to analyze 3-year patient survival following neutrophil engraftment using the lifelines (v0.24) package for the Python programming language (DOI:10.5281/zenodo.3677104).

$$h\left( t \right)=h_{0}\left( t \right)exp(b_{1}X_{1}+b_{2}X_{2}+\ldots+b_{n}X_{n} )$$

where h(t) is the expected hazard at time t, h_0_(t) is the baseline hazard. As predictors, *X*, we included the median total white blood cell count during the first 100 days following the time when FMT was usually performed (i.e. 37 days after neutrophil engraftment), sex, and age removing patients whothey lacked sufficient blood data during this post-engraftment period. Disease and stem cell graft source were used to stratify patients (using the “strata” parameter of the Cox proportional hazard fit function), as including them in the form of intercept terms violated the proportional hazards assumptions. This showed a mild but detectable association of total white blood cell counts during the investigated interval with improved survival:

|  | N: | 2,013 |  |  |  |
| --- | --- | --- | --- | --- | --- |
|  | events: | 668 |  |  |  |
|  | strata: | 'disease', 'hct-source' | |  |  |
|  | coef | exp(coef) | lower 0.95 | upper 0.95 | p-value |
| average total wbc count (z-scored) | -0.09 | 0.91 | -0.18 | 0 | 0.04 |
| age (z-scored) | 0.19 | 1.21 | 0.1 | 0.28 | <0.005 |
| SEX:F | 0.03 | 1.03 | -0.12 | 0.19 | 0.68 |

**Dynamic systems analyses**

*Covariates included in interval data*

For a given daily interval, a medication was considered present for at least part of the interval when it was administered on either endpoint. Administration events were obtained from parsing the institutional task data base which contains drug and treatment administrations performed on patients. For the microbiota, data from the end day were considered for the interval, and for blood stream infection data, either endpoint was considered. Homeostatic feedback calculations used the geometric mean of the white blood cell counts between the two endpoints. We included only those covariates that were present during at least 10 intervals.

Our data comprises >1.6M recorded administrations of 806 different medications with durations provided. All patients analyzed in stage 2 had available medication records, but ~10% of the patients without microbiome data had missing medication records and/or incomplete metadata. In case of missing continuous metadata, missing values were filled with means.

The stem cell graft source is a major determinant of engraftment times and can affect recovery dynamics^1,2^, and we therefore included intercept terms for unmodified peripheral blood stem cell grafts (PBSC), bone marrow (BM), T-cell depleted graft (*ex‑vivo*) by CD34+selection (TCD) and cord blood (cord) in stage 1.

Patients received a variety of conditioning regimens comprised of various doses of chemotherapy and, in some cases, irradiation. There are dozens of conditioning regimes for HCT^3^. The standard approach in the allo-HCT field for observational studies is to categorize them by the Bacigalupo classification, which uses three graded categories from most to least intense^4^: Myeloablative (“ABLATIVE”), Reduced Intensity (“REDUCE”), and Nonmyeloablative (“NONABL”). We included the conditioning intensity as indicator variables.

*Data exclusion for white blood cell dynamic (stage 1 and 2)*

Our dynamics analyses focus on the daily changes in white blood cell counts from one day to another during recovery of the circulatory immune cell system. To analyze the kinetics of this reconstitution, we excluded data when (see also flow chart below):

- On the day of HCT, patients were younger than 18 years
- Patients did not engraft
- Patients had a second transplant within 100 days
- Samples were taken outside the window of 100 days starting from neutrophil engraftment
- In case there were multiple blood samples per patient and day, the one closest to noon was chosen
- A patient died within the first 3 months after HCT
- A sample was taken within 1 week of FMT

This data exclusion is encoded as two separate columns in the tidy data table WBC.csv, by a Boolean indicator column named “include”, and the column “exclude_reason” of string type.

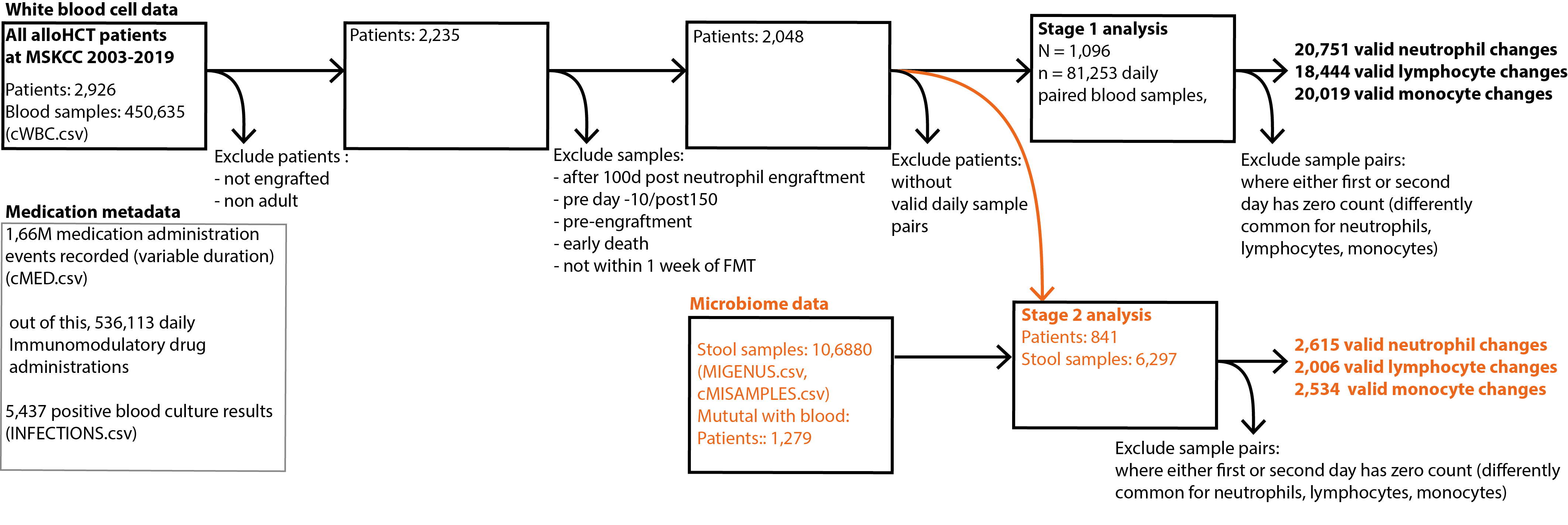

*Shotgun sequencing data processing*

We removed normal optical duplicates in paired FASTQ files using the clumpify.sh tool from the BBMap package (BBMap – Bushnell B. – <https://www.sourceforge.net/projects/bbmap/>), producing a pair of deduped read files. Using the bbduk.sh script in the BBMap package, we trimmed the right and left side of a read in a pair to Q10 using the Phred algorithm. A pair of reads was dropped if any one of them has a length shorter than 51 nucleotides after trimming. We trim 3’-end adapters using a kmer of length 31, and a shorter kmer of 9 at the other end of the read. One mismatch was allowed in this process, and we allowed adapter trimming based on pair overlap detection (which does not require known adapter sequences) using the ‘tbo’ parameter. We used the ‘tpe’ parameter to trim the pair of reads to the same length.

Removal of human contamination was done using Kneaddata with paired-end reads, employing BMTagger. The BMTagger database was built with human genome assembly GRCh38. After decontamination, the paired-end reads were concatenated to a single FASTQ file as the input for functional profiling with the Humann2 pipeline (main methods).

*Rationale for using estimations of total bacterial abundances for bi-directional analysis of bacterial and white blood cell dynamics*

In order to assess the association of bacterial populations with antibiotics, the ecosystem of other bacteria and the count of white blood cells in circulation, we employed a similar approach to that used in stage 1. To calculate the dependent variable, i.e. the log-changes per day in total counts, we had to estimate the absolute abundances of bacteria. This is needed because inferences of dynamic equations like the ones used here cannot be conducted on observations of changes in relative abundances. Briefly, a positive change in relative abundances of a bacterium could be due to the focal bacterium’s counts increasing, that of other species’ decreasing, both focal and others increasing or decreasing but the focal species faster or slower so, respectively. This means that when attempting to use observations of relative abundance changes as dependent variables to infer interactions like we do here, one would try to solve one more unknown than one has information for. Unless specific assumptions can be justified, one therefore must estimate a total count. We here do this by estimating the total population density using qPCR.

This is important when estimating the sign and the effect sizes of antibiotics, other bacterial species in the ecosystem or the white blood cells in circulation on the changes of a focal taxon. We do not know what aspects of the microbiota can be affected by the immune system, and it is entirely possible that the immune system evolved to “control” relative abundances to achieve microbiota homeostasis^5^. Yet, in order to estimate in which direction the immune system’s effect operates on any one taxon, for the reasons above, we require observations of total changes per taxon.

*Duke data collection*

Data at Duke university was collected analogously to the procedures at MSKCC. Complete blood counts were assessed on the Sysmex XN, Pro00050975 platform. Clinical metadata was obtained from the Duke Enterprise Data Unified Content Explorer (DEDUCE) and Duke Adult Blood and Marrow Transplant (ABMT) data base. All sequencing was conducted at MSKCC following the same protocols.
